# Efa6 protects axons and regulates their growth and branching by inhibiting microtubule polymerisation at the cortex

**DOI:** 10.1101/385658

**Authors:** Yue Qu, Ines Hahn, Meredith Lees, Jill Parkin, André Voelzmann, Karel Dorey, Alex Rathbone, Claire Friel, Viki Allan, Pilar Okenve-Ramos, Natalia Sanchez-Soriano, Andreas Prokop

## Abstract

Cortical collapse factors affect microtubule (MT) dynamics at the plasma membrane. They play important roles in neurons, as suggested by inhibition of axon growth and regeneration through the Arf activator Efa6 in *C. elegans*, and by neurodevelopmental disorders linked to the mammalian kinesin Kif21A. How cortical collapse factors influence axon growth is little understood. Here we studied them, focussing on the function of *Drosophila* Efa6 in experimentally and genetically amenable fly neurons. First, we show that *Drosophila* Efa6 can inhibit MTs directly without interacting molecules via an N-terminal 18 amino acid motif (MT elimination domain/MTED) that binds tubulin and inhibits microtubule growth *in vitro* and cells. If N-terminal MTED-containing fragments are in the cytoplasm they abolish entire microtubule networks of mouse fibroblasts and whole axons of fly neurons. Full-length Efa6 is membrane-attached, hence primarily blocks MTs in the periphery of fibroblasts, and explorative MTs that have left axonal bundles in neurons. Accordingly, loss of Efa6 causes an increase of explorative MTs: in growth cones, they enhance axon growth, in axon shafts, explorative MTs cause excessive branching, as well as atrophy through perturbations of MT bundles. Efa6 over-expression causes the opposite phenotypes. Taken together, our work conceptually links molecular and sub-cellular functions of cortical collapse factors to axon growth regulation and reveals new roles in axon branching and in the prevention of axonal atrophy. Furthermore, the MTED delivers a promising tool that can be used to inhibit MTs in a compartmentalised fashion when fusing it to specifically localising protein domains.

**Summary statement:** The cortical collapse factor Efa6 inhibits microtubule polymerising outside axonal bundles. Thereby it limits axon growth and branching, but preserves microtubule bundle organisation crucial for axon maintenance.

## Introduction

Axons are the cable-like neuronal extensions that wire the nervous system. They are only 0.1-15µm in diameter (Hoffman, 1995), but can be up to a meter long in humans (Debanne et al., 2011; Prokop, 2013a). It is a fascinating challenge to understand how axons can extend over these enormous distances and branch in orderly manners, but also how these delicate structures can be maintained for a lifetime, i.e. many decades in humans. It is not surprising that we gradually lose about half of our axons towards old age (Calkins, 2013; Marner et al., 2003), and that axon decay is a prominent neurodegenerative phenomenon (Adalbert and Coleman, 2012; Fang and Bonini, 2012; Medana and Esiri, 2003; Wang et al., 2012).

Essential for axon biology are the parallel bundles of microtubules (MTs) running all along the axon shaft; these bundles provide (1) structural support, (2) highways for life-sustaining cargo transport, and (3) a source of MTs that can leave these bundles to drive morphogenetic changes. Through being organised in this way, MTs essentially drive processes of axon growth, branching and maintenance (Conde and Caceres, 2009; Dent et al., 2011; Hahn et al., 2019; Prokop, 2013a; Voelzmann et al., 2016a). The dynamics of MTs are orchestrated through MT-binding and -regulating proteins, for most of which we know the molecular mechanisms of function. However, such knowledge alone is usually not sufficient to explain their cellular roles.

For example, cortical collapse factors are cell surface-associated proteins specifically inhibit MTs that approach the cell periphery. Previous reports suggested important roles for cortical collapse factors in regulating axon growth: the ARF activator Efa6 in *C. elegans* negatively impacts on developmental and regenerative axon growth (Chen et al., 2015; Chen et al., 2011; O’Rourke et al., 2010); the mammalian type 4 kinesin KIF21A also affects axon growth and links to the neurodevelopmental eye movement disorder “congenital fibrosis of extraocular muscles” (OMIM reference <u>#135700</u>; Heidary et al., 2008; Tiab et al., 2004; van der Vaart et al., 2013). However, we can currently only hypothesise how the molecular function of these two collapse factors links to axon growth, most likely by acting in growth cones (GCs).

GCs are the amoeboid tip structures where axons extend to wire the nervous system during development or regeneration. The axonal MT bundles terminate in the centre of GCs; from here, single MTs splay into the actin-rich periphery of GCs. These explorative MTs can trigger extension of the entire MT bundle into their direction, thus elongating the axon (Dent et al., 2011; Lowery and van Vactor, 2009; Prokop et al., 2013); their (partial) inhibition through cortical collapse factors could provide a potential mechanism through which cortical collapse factors negatively impact on axon growth.

In line with this argumentation, and depending on where cortical collapse factors are present and functionally active, further functional predictions could be made: for example, collateral branching of axons along their shafts has been described to depend on explorative MTs that leave the parallel axonal bundles and polymerise towards the periphery (Kalil and Dent, 2014; Lewis et al., 2013; Tymanskyj et al., 2017; Yu et al., 2008). Cortical collapse factors might therefore be negative regulators of axon branching.

Other roles might concern axon maintenance: the model of ‘local axon homeostasis’ states that the force-enriched environment in axons biases MTs to buckle or project out of the bundle to seed pathological areas of MT disorganisation (Hahn et al., 2019; Prokop, 2016; Voelzmann et al., 2016a). By inhibiting off-track MTs in the axon shaft, cortical collapse factors might prevent such processes, acting in parallel to other bundle-maintaining factors. For example, spectraplakins serve as spacers that keep polymerising MTs away from the cortex by linking the tips of extending MTs to the axonal surface and guiding them into parallel bundles (Alves-Silva et al., 2012). Their deficiency in any organism causes severe MT disorganisation, potentially explaining human dystonin-linked HSAN6 (’type 6 hereditary sensory and autonomic neuropathy’’; <u>#614653</u>; Voelzmann et al., 2017). If our hypothesis is correct, loss of cortical collapse factors in axon shafts would also cause MT disorganisation, but through a very different mechanistic route.

Here we make use of *Drosophila* neurons as a well-established, powerful model for studying roles of MT regulators (Hahn et al., 2019; Prokop et al., 2013; Sánchez-Soriano et al., 2007). Using *in vitro* and cellular assays, we show that *Drosophila* Efa6 is a cortical collapse factor acting through its N-terminal MT-eliminating domain (MTED). We find that the MTED binds tubulin and blocks MT polymerisation *in vitro* which shows that the effect of the peptide is due to a direct interaction between the peptide and tubulin and does not require any other molecules. By localising to neuronal membranes, it only abolishes explorative MTs. This subcellular role translates into negative regulation of axon growth and branching and the prevention of pathological MT disorganisation, both in cultured neurons and *in vivo*. We propose Efa6 to function as a quality control or axonal maintenance factor that keeps explorative MTs in check, thus playing a complementary role to spectraplakins that prevent MTs from leaving axonal bundles.

## Methods

### Fly stocks

Loss-of-function mutant stocks used in this study were the two deficiencies uncovering the *Efa6* locus *Df(3R)Exel6273* (94B2-94B11 or 3R:22,530,780..22,530,780) and *Df(3R)ED6091i* (94B5-94C4 or 3R:22,587,681..22,587,681), *shot^3^* (the strongest available allele of *short stop*) (Kolodziej et al., 1995; Sánchez-Soriano et al., 2009), and the null mutant alleles *Efa6^KO#1^*, *Efa6^GX6[w-]^*, *Efa6^GX6[w+]^* and *Arf51F^GX16[w-]^* (all genomically engineered precise deletions) (Huang et al., 2009). Gal4 driver lines used were the pan-neuronal lines *sca-Gal4* (strongest in embryos) (Sánchez-Soriano et al., 2010) and *elav-Gal4* (1^st^ and 3^rd^ chromosomal, both expressing at all stages) (Luo et al., 1994), as well as the *ato-Gal4* line expressing in a subset of neurons in the adult brain (Hassan et al., 2000; Voelzmann et al., 2016b). Lines for targeted gene expression were *UAS-Efa6^RNAi^* (VDRC #42321), *UAS-Gal80^ts^*(Zeidler et al., 2004), *UAS-Eb1-GFP* (Alves-Silva et al., 2012), *UAS-α-tubulin84B* (Grieder et al., 2000) and *UAS-tdTomato* (Zschätzsch et al., 2014). Efa6 expression was detected via the genomically engineered *Efa6-GFP* allele, where a GFP was inserted after the last amino acid in exon 14 (Huang et al., 2009).

### *Drosophila* primary cell culture

*Drosophila* primary neuron cultures were performed as published previously (Prokop et al., 2012; Qu et al., 2017). In brief, stage 11 embryos were treated for 1 min with bleach to remove the chorion, sterilized for ∼30 s in 70% ethanol, washed in sterile Schneider’s/FCS, and eventually homogenized with micro-pestles in 1.5 centrifuge tubes containing 21 embryos per 100 μl dispersion medium (Prokop et al., 2012) and left to incubated for 5 min at 37°C. Cells are washed with Schneider’s medium (Gibco), spun down for 4 mins at 650 g, supernatant was removed and cells re-suspended in 90 µl of Schneider’s medium containing 20% fetal calf serum (Gibco). 30 μl drops were placed on cover slips. Cells were allowed to adhere for 90-120 min either directly on glass or on cover slips coated with a 5 µg/ml solution of concanavalin A, and then grown as a hanging drop culture for hours or days at 26°C as indicated.

To abolish maternal rescue of mutants, i.e. masking of the mutant phenotype caused by deposition of normal gene product from the healthy gene copy of the heterozygous mothers in the oocyte (Prokop, 2013b), we used a pre-culture strategy (Prokop et al., 2012; Sánchez-Soriano et al., 2010) where cells were kept for 5 days in a tube before they were plated on a coverslip.

For the transfection of *Drosophila* primary neurons, a quantity of 70-75 embryos per 100 μl dispersion medium was used. After the washing step and centrifugation, cells were re-suspended in 100 μl transfection medium [final media containing 0.1-0.5 μg DNA and 2 μl Lipofecatmine 2000 (L2000)]. To generate this media, dilutions of 0.1-0.5 μg DNA in 50 μl Schneider’s medium and 2 μl L2000 in 50 μl Schneider’s medium were prepared, then mixed together and incubated at room temperature for 5-30 mins, before being added to the cells in centrifuge tubes where they were kept for 24 hrs at 26°C. Cells were then treated again with dispersion medium, re-suspended in culture medium and plated out as described above.

For temporally controlled knock-down experiments we used flies carrying the driver construct *elav-Gal4*, the knock-down construct *UAS-Efa6-RNAi*, and the temperature-sensitive Gal4 inhibitor *UAS-Gal80^ts^*, all in parallel. At the restrictive temperature of 19°C, Gal80^ts^ blocks Gal4-induced expression of *Efa6-RNAi*, and this repressive action is removed at the permissive temperature of 27°C where Gal80^ts^ is non-functional. Control neurons were from flies carrying only the *Gal4/Gal80* (control 1 in Fig. 8K) or only the *Efa6-RNAi* transgene (control 2).

### Fibroblast cell culture

NIH/3T3 fibroblasts were grown in DMEM supplemented with 1% glutamine (Invitrogen), 1% penicillin/streptomycin (Invitrogen) and 10% FCS in culture dishes (100 mm with vents; Fisher Scientific UK Ltd) at 37°C in a humidified incubator at 5% CO_2_. Cells were split every 2-3 d, washed with pre-warmed PBS, incubated with 4 ml of Trypsin-EDTA (T-E) at 37°C for 5 min, then suspended in 7 ml of fresh culture medium and eventually diluted (1/3-1/20 dilution) in a culture dish containing 10 ml culture media.

For transfection of NIH/3T3 cells, 2 ml cell solution (∼10^5^ cells per ml) were first transferred to 6-well plates, and grown overnight to double cell density. 2 µg of DNA and 2 µl Plus reagent (Invitrogen) were added to 1 ml serum-free media in a centrifuge tube, incubated for 5 mins at RT, then 6 µl Lipofectamine (Invitrogen) were added, and incubated at RT for 25 mins. Cells in the 6-well plate were washed with serum-free medium and 25 mins later DNA/Lipofectamine was mixed into the medium (1/1 dilution). Plates were incubated for 3 hrs at 37°C, washed with 2 ml PBS, 400 µl trypsin were added for 5 mins (37°C), then 3 ml complete medium; cells were suspended and added in 1 ml aliquots to 35 mm glass-bottom dishes (MatTek) coated with fibronectin [300 µl of 5 µg/ml fibronectin (Sigma-Aldrich) placed in the center of a MatTek dish for 1 hr at 37°C, then washed with PBS]; 1 ml of medium was added and cells grown for 6 hrs or 24 hrs at 37°C in a CO_2_ incubator. For live imaging, the medium was replaced with 2 ml Ham’s F-12 medium + 4% FCS.

### Dissection of adult brains

To analyse the function of Efa6 in MT bundle integrity in medulla axons *in vivo*, flies were aged at 29°C. Flies were maintain in groups of up to 20 flies of the same gender (Stefana et al., 2017) and changed into new tubes every 3-4 days. Brain dissections were performed in Dulbecco’s PBS (Sigma, RNBF2227) after briefly sedating them on ice. Dissected brains with their laminas and eyes attached were placed into a drop of Dulbecco’s PBS on MatTek glass bottom dishes (P35G1.5-14C), covered by coverslips and immediately imaged with a 3i Marianas Spinning Disk Confocal Microscope.

To measure branching in *ato-Gal4 Drosophila* neurons, adult brains were dissected in Dulbecco’s PBS and fixed with 4% PFA for 15 min. Antibody staining and washes were performed with PBS supplemented with 0.3% Triton X-100. Specimens were embedded in Vectashield (VectorLabs).

### Immunohistochemistry

Primary fly neurons and fibroblasts were fixed in 4% paraformaldehyde (PFA) in 0.05 M phosphate buffer (PB; pH 7–7.2) for 30 min at room temperature (RT); for anti-Eb1 staining, ice-cold +TIP fix (90% methanol, 3% formaldehyde, 5 mM sodium carbonate, pH 9; stored at −80°C and added to the cells) (Rogers et al., 2002) was added for 10 mins. Adult brains were dissected out of their head case in PBS and fixed with 4% PFA in PBS for 1 hr, followed by a 1hr wash in PBT.

Antibody staining and washes were performed with PBT. Staining reagents: anti-tubulin (clone DM1A, mouse, 1:1000, Sigma; alternatively, clone YL1/2, rat, 1:500, Millipore Bioscience Research Reagents); anti-DmEb1 (gift from H. Ohkura; rabbit, 1:2000) (Elliott et al., 2005); anti-Elav (mouse, 1:1000, DHB); anti-GFP (goat, 1:500, Abcam); Cy3-conjugated anti-HRP (goat, 1:100, Jackson ImmunoResearch); F-actin was stained with Phalloidin conjugated with TRITC/Alexa647, FITC or Atto647N (1:100 or 1:500; Invitrogen and Sigma). Specimens were embedded in ProLong Gold Antifade Mountant.

### Microscopy and data analysis

Standard documentation was performed with AxioCam monochrome digital cameras (Carl Zeiss Ltd.) mounted on BX50WI or BX51 Olympus compound fluorescent microscopes. For the analysis of *Drosophila* primary neurons, we used two well established parameters (Alves-Silva et al., 2012; Sánchez-Soriano et al., 2010): axon length (from cell body to growth cone tip; measured using the segmented line tool of ImageJ) and the degree of MT disorganization in axons which was either measured as binary score or ratio (percentage of neurons showing obvious MT disorganisation in their axons) or as “MT disorganisation index” (MDI) (Qu et al., 2017): the area of disorganisation was measured using the freehand selection in ImageJ; this value was then divided by axon length (see above) multiplied by 0.5 μm (typical axon diameter, thus approximating the expected area of the axon if it were not disorganised). For Eb1::GFP comet counts, neurons were subdivided into axon shaft and growth cones (GC): the proximal GC border was set where the axon widens up (broader GCs) or where filopodia density increases significantly (narrow GCs). MT loss in fibroblasts was assessed on randomly chosen images of successfully transfected, GFP-expressing fibroblasts, stained for tubulin and actin. Images were derived from at least 2 independent experimental repeats performed on different days, for each of which at least 3 independent culture wells were analysed by taking a minimum of 20 images per well. Due to major differences in plasma membrane versus cytoplasmic localisation of constructs, their expression strengths could not be standardised. Assuming a comparable expression strength distribution, we therefore analyse all transfected cells in the images and assigned them to three categories: MTs intact, damaged or gone (Fig. 3G-G’’). To avoid bias, image analyses were performed blindly, i.e. the genotype or treatment of specimens was masked. To analyse ruffle formation in fibroblasts, cells were stained with actin and classified (with or without ruffles).

To assess the degree of branching, we measured axonal projections of dorsal cluster neurons in the medulla, which is part of the optic lobe in the adult brain (Hassan et al., 2000; Voelzmann et al., 2016b). These neurons were labelled by expressing *UAS-myr-tdTomato* via the *ato-Gal4* driver either alone (control), together with *UAS-Efa6^RNAi^*or together with *UAS-Efa6-FL-GFP*. We analysed them in young brains (2-5 d after eclosure of flies from their pupal case) or old brains (15-18 d). Z-stacks of adult fly brains (optic lobe area) were taken with a Leica DM6000 B microscope and extracted with Leica MM AF Premier software. They were imaged from anterior and the number of branches was quantified manually. Branches were defined as the protrusions from the DC neuron axons in the medulla. Branches in fly primary neurons at 5DIV were also counted manually and defined as MT protrusions from main axon.

To measure MT disorganisation in the optic lobe of adult flies, *GMR31F10-Gal4* (Bloomington #49685) was used to express *UAS-α-tubulin84B-GFP* (Grieder et al., 2000) in a subset of lamina axons which projects within well-ordered medulla columns (Prokop and Meinertzhagen, 2006). Flies were left to age for 26-27 days (about half their life expectancy) and then their brains were dissected out, mounted in Mattek dishes and imaged using a 3i spinning disk confocal system at the ITM Biomedecial imaging facility at the University of Liverpool. A section of the medulla columns comprising the 4 most proximal axonal terminals was used to quantify the number of swellings and regions with disorganised MTs.

Time lapse imaging of cultured primary neurons (in Schneider’s/FCS) and fibroblasts (in Ham’s F-12/FCS) was performed on a Delta Vision Core (Applied Precision) restoration microscope using a [*100x/1.40 UPlan SAPO (Oil)*] objective and the Sedat Quad filter set (*Chroma #89000*). Images were collected using a Coolsnap HQ2 (Photometrics) camera. The temperature was set to 26°C for fly neurons and 37°C for fibroblasts. Time lapse movies were constructed from images taken every 2 s for 2 mins. To analyse MT dynamics, Eb1::GFP comets were tracked manually using the “manual tracking” plug-in of ImageJ.

For statistical analyses, Kruskal–Wallis one-way ANOVA with *post hoc* Dunn’s test or Mann–Whitney Rank Sum Tests (indicated as P_MW_) were used to compare groups, and χ2 tests (indicated as P_X2_) were used to compare percentages. All raw data of our analyses are provided as supplementary Excel/Prism files.

### Molecular biology

EGFP tags are based on pcDNA3-EGFP or pUAST-EGFP. All *Drosophila melanogaster* efa6 constructs are based on cDNA cloneIP15395 (Uniprot isoform C, intron removed). *Caenorhabditis elegans* efa-6 (Y55D9A.1a) constructs are derived from pCZGY1125-efa-6-pcr8 (kindly provided by Andrew Chisholm). *Homo sapiens* PSD1 (ENST00000406432.5, isoform 202) constructs were PCR-amplified from pLC32-hu-psd1-pcr8 vector (kindly provided by Andrew Chisholm). *Homo sapiens* PSD2 (ENST00000274710.3, isoform 201, 771aa) constructs were PCR-amplified from pLC33-hu-psd2-pcr8 vector (kindly provided by Andrew Chisholm). *Homo sapiens* PSD3 was PCR-amplified from pLC34 hu-psd3-pcr8 vector (kindly provided Andrew Chisholm). Note that the PSD3 cDNA clone is most closely related to isoform 201 (ENST00000286485.12: 513aa) and therefore lacks the putative N-terminus found in isoform 202 (ENST00000327040.12). However, the putative MTED core sequence is encoded in the C-terminal PH domain (Fig.2C), not the potential N-terminus. *Homo sapiens* PSD4 (ENST00000441564.7, isoform 205) was PCR-amplified from pLC35-hu-psd4-pcr8 vector (kindly provided by Andrew Chisholm).The CAAX motif is derived from human KRAS. The *Dm*Efa6-NtermΔSxiP::EGFP (aa1-410) insert was synthesised by GeneArt Express (ThermoFisher). All construct were cloned using standard (SOE) PCR/ligation based methods, and constructs and inserts are detailed in Table T1. To generate transgenic fly lines, *P*[*acman*]*M-6-attB-UAS-1-3-4* constructs were integrated into *PBac{yellow*[*+*]*-attP-3B}VK00031* (Bloomington line #9748) via PhiC31 mediated recombination (outsourced to Bestgene Inc.).

**Tab. T1.** co=codon optimised; Dm=Drosphila melanogaster; Ce=Caenorhabditis elegans; Hs=Homo sapiens

| final vector | Source | insert |
| --- | --- | --- |
| **pcDNA3-EGFP** | Addgene | <u>*XhoI*-EGFP-*XbaI*</u> |
| **pUAST-*Ascl*-*PacI*-EGFP** | this study | <u>*KpnI*, *Ascl*, *PacI*-EGFP-*XbaI*</u> |
| **pUAST-*DmEfa6*FL-EGFP (aa1-1387)** | this study | <u>*KpnI*, *Ascl*-**kozak**-*Efa6* (aa1-1387)-**GSGSGS-EGFP**-*PacI*, *XbaI*</u> |
| ***P[acman]M-6-attB-UAS-1-3-4-DmEfa6*FL-EGFP (aa1-1387)** | this study | <u>*Ascl*-**kozak**-*DmEfa6* (aa1-1387)-**GSGSGS-EGFP**-*PacI*</u> |
| **pcDNA3.1-*DmEfa6*FL-EGFP (aa1-1387)** | this study | <u>*KpnI*, *Ascl*-**kozak**-*DmEfa6* (aa1-1387)-**GSGSGS-EGFP**-*PacI*, *XbaI*</u> |
| **pUAST-*DmEfa6*ΔCterm-EGFP (aa1-894)** | this study | <u>*KpnI*, *Ascl*-**kozak**-*DmEfa6*ΔCterm (aa1-894)-**GSGSGS-EGFP**-*PacI*, *XbaI*</u> |
| ***P[acman]M-6-attB-UAS-1-3-4-DmEfa6\DeltaCterm-EGFP (aa1-894)*** | this study | <u>*Ascl*-**kozak**-*DmEfa6*ΔCterm (aa1-894)-**GSGSGS-EGFP**-*PacI*</u> |
| **pcDNA3.1-*DmEfa6*ΔCterm-EGFP (aa1-894)** | this study | <u>*KpnI*, *Ascl*-**kozak**-*DmEfa6*ΔCterm (aa1-894)-**GSGSGS-EGFP**-*PacI*, *XbaI*</u> |
| **pUAST-*DmEfa6*-Nterm-EGFP (aa1-410)** | this study | <u>*KpnI*, *Ascl*-**kozak**-*DmEfa6*-Nterm (aa1-410)-**GSGSGS-EGFP**-*PacI*, *XbaI*</u> |
| ***P[acman]M-6-attB-UAS-1-3-4-DmEfa6*-Nterm-EGFP (aa1-410)** | this study | <u>*Ascl*-**kozak**-*DmEfa6*-Nterm (aa1-410)-**GSGSGS-EGFP**-*PacI*</u> |
| **pcDNA3.1-*DmEfa6*-Nterm-EGFP (aa1-410)** | this study | <u>*KpnI*, *Ascl*-**kozak**-*DmEfa6*-Nterm (aa1-410)-**GSGSGS-EGFP**-*PacI*, *XbaI*</u> |
| **pUAST-*DmEfa6*-Nterm-CAAX-EGFP (aa1-410)** | this study | <u>*KpnI*, *Ascl*-**kozak**-*DmEfa6*-Nterm (aa1-410)-**GSGSGS-EGFP-CAAX[*KRAS*]**-*PacI*, *XbaI*</u> |
| ***P[acman]M-6-attB-UAS-1-3-4-DmEfa6*-Nterm-CAAX-EGFP (aa1-410)** | this study | <u>*Ascl*-**kozak**-*DmEfa6*-Nterm (aa1-410)-**GSGSGS-EGFP-CAAX[*KRAS*]**-*PacI*</u> |
| **pcDNA3.1-*DmEfa6*-Nterm-CAAX-EGFP (aa1-410)** | this study | <u>*KpnI*, *Ascl*-**kozak**-*DmEfa6*-Nterm (aa1-410)-**GSGSGS-EGFP-CAAX[*KRAS*]**-*PacI*, *XbaI*</u> |
| **pUAST-DmEfa6-NtermΔSxiP-EGFP (aa1-410)** | this study | <u>KpnI, AscI-kozak-DmEfa6-NtermΔSxiP (aa1-410; SQIP>AAAA; SRIP>AAAA)-GSGSGS-EGFP-PacI, XbaI</u> |
| **pcDNA3.1-DmEfa6-NtermΔSxiP-EGFP (aa1-410)** | this study | <u>KpnI, AscI-kozak-DmEfa6-NtermΔSxiP (aa1-410; SQIP>AAAA; SRIP>AAAA)-GSGSGS-EGFP-PacI, XbaI</u> |
| **pUAST-DmEfa6-NtermΔMTED-EGFP (aa1-300)** | this study | <u>KpnI, AscI-kozak-DmEfa6-NtermΔMTED (aa1-300)-GSGSGS-EGFP-PacI, XbaI</u> |
| **pcDNA3.1-DmEfa6-NtermΔMTED-EGFP (aa1-300)** | this study | <u>KpnI, AscI-kozak-DmEfa6-NtermΔMTED (aa1-300)-GSGSGS-EGFP-PacI, XbaI</u> |
| **pUAST-DmEfa6ΔNerm-EGFP (aa851-1387)** | this study | <u>KpnI, AscI-kozak-DmEfa6ΔNerm (aa851-1387)-GSGSGS-EGFP-PacI, XbaI</u> |
| **pcDNA3.1-DmEfa6ΔNerm-EGFP (aa851-1387)** | this study | <u>KpnI, AscI-kozak-DmEfa6ΔNerm (aa851-1387)-GSGSGS-EGFP-PacI, XbaI</u> |
| **pUAST-DmEfa6-MTED-EGFP (aa322-341)** | this study | <u>KpnI, AscI-kozak-DmEfa6-MTED (aa322-341)-GSGSGS-EGFP-PacI, XbaI</u> |
| **pcDNA3.1-DmEfa6-MTED-EGFP (aa322-341)** | this study | <u>KpnI, AscI-kozak-DmEfa6-MTED (aa322-341)-GSGSGS-EGFP-PacI, XbaI</u> |
| **pcDNA3.1-CeEfa6-FL-EGFP (aa1-816)** | this study | <u>KpnI, AscI-kozak-CeEfa6 (aa1-816)-GSGSGS-EGFP-PacI, XbaI</u> |
| **pcDNA3.1-CeEfa6-Nterm-EGFP (aa1-152)** | this study | <u>KpnI, AscI-kozak-CeEfa6-Nterm (aa1-152)-GSGSGS-EGFP-PacI, XbaI</u> |
| **pcDNA3.1-CeEfa6-MTED-EGFP (aa24-42)** | this study | <u>KpnI, AscI-kozak-CeEfa6-MTED (aa24-42)-GSGSGS-EGFP-PacI, XbaI</u> |
| **pcDNA3.1-HsPSD1-FL-EGFP (aa1-1024)** | this study | <u>KpnI, AscI-kozak-HsPSD1 (aa1-1024)-GSGSGS-NotI-EGFP-PacI, XbaI</u> |
| **pcDNA3.1-HsPSD1-Nterm-EGFP (aa1-280)** | this study | <u>KpnI, AscI-kozak-HsPSD1-Nterm (aa1-280)-GSGSGS-NotI-EGFP-PacI, XbaI</u> |
| **pcDNA3.1-HsPSD1-MTED-EGFP (aa31-49)** | this study | <u>KpnI, AscI-kozak-HsPSD1-MTED (aa31-49)-GSGSGS-NotI-EGFP-PacI, XbaI</u> |
| **pcDNA3.1-HsPSD2-FL-EGFP (aa1-771)** | this study | <u>KpnI, AscI-kozak-HsPSD2 (aa1-771)-GSGSGS-NotI-EGFP-PacI, XbaI</u> |
| **pcDNA3.1-HsPSD3-EGFP (aa515-1047)** | this study | <u>KpnI, AscI-kozak-HsPSD3 (aa515-1047)-GSGSGS-NotI-EGFP-PacI, XbaI</u> |
| **pcDNA3.1-HsPSD4-FL-EGFP (aa1-1027)** | this study | <u>KpnI, AscI-kozak-HsPSD4 (aa1-1027)-GSGSGS-NotI-EGFP-PacI, XbaI</u> |
| **pcDNA3.1-co-HsPSD1-MTED-EGFP (aa31-49)** | this study | <u>KpnI, AscI-kozak-HsPSD1-MTED (aa31-49)-GSGSGS-EGFP-PacI, XbaI</u> |
| **pcDNA3.1-co-CeEfa6-MTED-EGFP (aa24-42)** | this study | <u>KpnI, AscI-kozak-CeEfa6-MTED (aa24-42)-GSGSGS-EGFP-PacI, XbaI</u> |
| **pcDNA3.1-co-DmEfa6-MTED-EGFP (aa322-341)** | this study | <u>KpnI, AscI-kozak-DmEfa6-MTED (aa322-341)-GSGSGS-EGFP-PacI, XbaI</u> |
| **pCS107-DmEfa6-Nterm-EGFP** | this study | <u>NotI-kozak-DmEfa6-Nterm (aa1-410)-GSGSGS-EGFP-StuI</u> |
| **RFP (Xenopus injection)** | tba | tba |
| **pFastBac-His6-MCAK-EGFP-StrepII** | tba | His6-MCAK::EGFP-StrepII |
| **pFastBac-His6-DmEfa6ΔCterm-EGFP-StrepII (aa1-894)** | this study | His6-DmEfa6ΔCterm::EGFP-StrepII (aa1-894) |

### *In silico* analyses

To generate the **phylogenetic tree** of Efa6/PSD full length isoforms and N-terms of different species (see Fig. S1), their amino acid sequences were aligned using Muscle or ClustalO (Goujon et al., 2010; McWilliam et al., 2013; Sievers et al., 2011). ProtTest (Abascal et al., 2005; Darriba et al., 2011) was used to determine amino acid frequencies in the protein datasets and to identify the optimal amino acid substitution model to be used for the Bayesian inference (VT+I+G+F). CUDA-Beagle-optimised MrBayes (Ronquist et al., 2012) was run using the VT+I+G+F model [prset statefreqpr=fixed(empirical); lset rates=invgamma] using 5 chains (1 heated) and 9 parallel runs until the runs converged and standard deviation of split frequencies were below 0.015 (0.06 for N-terms); PSRF+ was 1.000 and min ESS was >1300 for the TL, alpha and pinvar parameters. The *Drosophila melanogaster* Sec7-PH domain-containing protein Steppke was used as outgroup in the full length tree. Archaeopteryx (Han and Zmasek, 2009) was used to depict the MrBayes consensus tree showing branch lengths (amino acid substitutions per site) and Bayesian posterior probabilities.

To **identify a potential MTED in PSD1**, previously identified Efa6 MTED motifs (O’Rourke et al., 2010) of 18 orthologues were aligned to derive an amino acid logo. Further orthologues were identified and used to refine the logo. Invariant sites and sites with restricted amino acid substitutions were determined (most prominently MxG-stretch). Stretches containing the invariant MxG stretch were aligned among vertebrate species to identify potential candidates. Berkley’s Weblogo server (Crooks et al., 2004) was used to generate amino acid sequence logos for each phylum using MTED (ExxxMxGE/D) and MTED-like (MxGE/D) amino acid stretches.

### *In vitro* analyses

#### Protein Expression and Purification

*Drosophila* Efa6-ΔCterm was cloned into a modified pFastBac vector containing an N-terminal *His6* tag and C-terminal *eGFP* and *StrepII* tags. Recombinant protein was expressed in Sf9 insect cells for 72 hours using a *Baculovirus* system. The protein was purified via a two-step protocol of Ni-affinity using a 1ml His-Trap column (GE Healthcare) in Ni-affinity buffer [50 mM Tris pH 7.5, 300 mM NaCl, 1mM Mg Cl_2_, 10 % (v/v) glycerol] and elution with 200mM imidazole, followed by Step-tag affinity chromatography using StepTactin resin (GE Healthcare) in BRB20, 75mM KCl. 0.1% Tween 20, 10% (v/v) glycerol and elution with 5mM desthiobiotin. MTED peptide (Genscript) was shipped as lyophilised powder with a purity of 95.2%. Upon arrival peptide was dissolved in ultrapure water and used directly.

#### MT binding assays

GMPCPP-stabilised, rhodamine-labeled MTs were adhered to the surface of flow chambers (Helenius et al., 2006). 20 nM Efa6-ΔCterm::GFP (in BRB20 pH 6.9, 75mM KCl, 0.05% Tween20, 0.1 mg/ml BSA, 1% 2-mercaptoethanol, 40mM glucose, 40 mg/ml glucose oxidase, 16 mg/ml catalase) or 20 nM MCAK::GFP (in the same buffer plus 1 mM ATP and 1 mM taxol) was introduced to the MT-containing channel. Images were recorded using a Zeiss Observer.Z1 microscope equipped with a Zeiss Laser TIRF 3 module, QuantEM 512SC EMCDD camera (Photometrics) and 100x objective (Zeiss, alphaPlanApo/1.46NA oil). Images of rhodamine-labeled MTs using a lamp as the excitation source and GFP fluorescence using TIRF illumination via a 488 nm laser were collected as described (Patel et al., 2016). For both rhodamine and GFP imaging an exposure time of 100 ms was used. The mean GFP intensity on individual MTs was determined from the mean pixel intensity of lines drawn along the long-axis of individual microtubules in Fiji (Schindelin et al., 2012). The rhodamine signal was used to locate the position of MTs in the GFP images. Intensity from a region of background was subtracted.

#### MT depolymerisation assays

GMPCPP-stabilised, rhodamine-labelled MTs were adhered to the surface of flow chambers (Helenius et al., 2006). Images of a field of fluorescent microtubules were recorded using a Zeiss Observer.Z1 microscope, collecting 1 image every 5 s with an exposure time of 100 ms. Efa6-ΔCterm::GFP (14 nM), MCAK (40 nM) in solution (BRB20 pH 6.9, 75mM KCl, 1mM ATP, 0.05% Tween 20, 0.1 mg/ml BSA, 1% 2-mercaptoethanol, 40mM glucose, 40 mg/ml glucose oxidase, 16 mg/ml catalase) were added to the channel 1 min after acquisition had commenced. Depolymerisation rates were determined from plots of the length of individual microtubules versus time, obtained by thresholding and particle analysis of images using Fiji (Schindelin et al., 2012).

#### *Xenopus* oocyte assays

cytosol extracts from *Xenopus* oocytes were obtained as described in (Allan and Vale, 1991). **MT depolymerisation** was assessed in a microscopic flow chamber (Vale and Toyoshima, 1988) where *Xenopus* cytosol (1 µl cytosol diluted with 20 µl acetate buffer) was incubated for 20 min to allow MTs to polymerise. Then cytosol was exchanged by flow through with Efa6-ΔCterm::GFP, MCAK or synthetic MTED peptide (all 20 nM in acetate buffer pH 7.4: 100 mM K-Acetate, 3 mM Mg-Acetate, 5 mM EGTA, 10 mM HEPES), and MT length changes observed by recording 10 random fields via VE-DIC microscopy (Allan, 1993; Allan and Vale, 1991). **MT polymerisation** was analysed in a microscope flow cell containing 9 µl diluted *Xenopus* cytosol (see above) to which 1 µl acetate buffer was added, either alone or containing 20nM MTED. After 10 min, 20 random fields were recorded via VE-DIC microscopy for each condition and the numbers of MTs per field counted.

For the *in vivo* assay, *Xenopus* embryos were injected in one blastomere at the 4-cell stage with 200 ng of mRNA encoding Efa6-Nterm::GFP or mCherry alone. The embryos were imaged at stage 10.25 (Heasman, 2006) with a Leica fluorescent stereoscope.

### Microtubule growth assays

30μM porcine brain tubulin (25% rhodamine-labelled) was incubated in 80mM PIPES pH6.9, 5mM MgCl_2_, 1mM EGTA, 5% DMSO and 1mM GTP at 37°C for 30min in the presence of either no peptide, 30μM MTED peptide or 300μM MTED peptide. The reactions were then diluted 60-fold into BRB80 buffer (80mM PIPES pH6.9, 1mM MgCl_2_, 1mM EGTA) containing 1mM taxol. Samples were added to channels constructed from poly-lysine coated cover glasses, washed with BRB80, 1mM taxol and imaged by fluorescence microscopy.

### Tubulin pull-down assays

MTED peptide was coupled to cyanogen bromide-activated Sepharose beads (GE Healthcare). 30μM porcine brain tubulin was incubated with either peptide-coated or uncoated Sepharose beads in BRB80, 0.2% Tween 20 for 30mins at 20°C. The beads were washed three times with a 2:1 *v/v* ratio of BRB80, 0.2% Tween 20 to beads. An equal volume of 2x Laemmli buffer was added to the washed beads, incubated at 90°C for 5min, spun down and supernatant run on a 12% SDS-PAGE gel.

## Results

### Efa6 is widely expressed in *Drosophila* neurons and restricts axonal growth

To evaluate the function of Efa6 in neurons, we first determined its expression in the nervous system. We used a genomically engineered fly line in which the endogenous *Efa6* gene was GFP-tagged (*Efa6-GFP*; Huang et al., 2009). These animals widely express Efa6::GFP throughout the CNS at larval and adult stages (Fig. 1F-I). We cultured primary neurons from this fly line to analyse the subcellular distribution of Efa6. In young neurons at 6 hrs *in vitro* (6HIV) and in mature neurons at 5 days *in vitro* (5DIV), Efa6 was localised throughout cell bodies and axons (Fig. 1B, E).

**Fig. 1.**
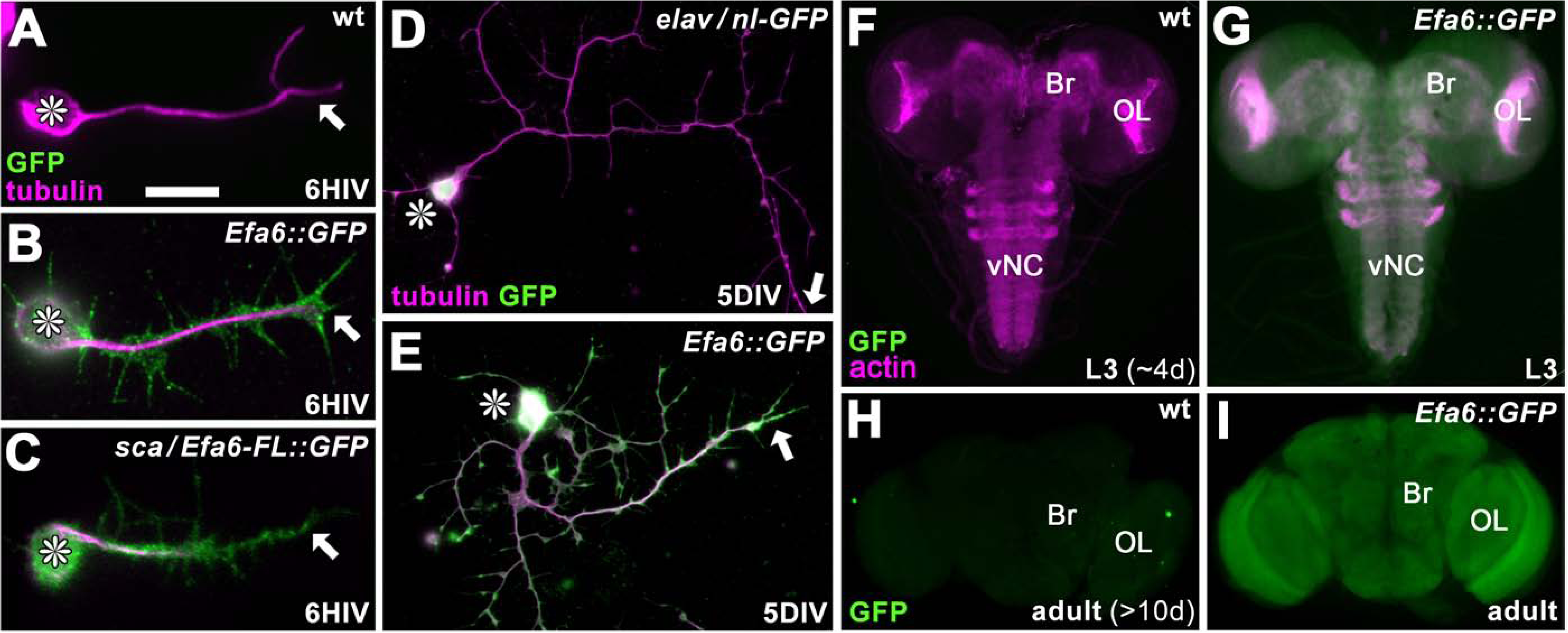
Efa6 is expressed throughout neurons at all developmental stages A-E) Images of primary *Drosophila* neurons at 6HIV or 5DIV (at indicated bottom right), stained for tubulin (magenta) and GFP (green); control neurons are wild-type (wt) or express *elav-Gal4*-driven nuclear GFP (*elav / nl-GFP*), whereas further neurons are either derived from the endogenously tagged Efa6::GFP line or express Efa6-FL::GFP under the control of sca-Gal4 (*sca / Efa6-FL::GFP*); asterisks indicate cell bodies and arrow the axon tips. F-I) Late larval CNSs at about 4d of development from egg lay (L3; F,G) and adult CNSs from 10d old flies (H,I) derived from control wild-type animals (wt) or the Efa6::GFP line (*Efa6::GFP*), stained for GFP and actin (Phalloidin, only larval preparations); OL, optic lobe; Br, central brain; vNC, ventral nerve cord. Scale bar in A represent 15µm in A-C, 25µm in D and E, 75µm in F and G, 130µm in H and I.

We next determined whether *Drosophila* Efa6 has an impact on axon growth, using fly lines with decreased or abolished Efa6 expression: Efa6 knock-down (*Efa6-RNAi*), overlapping deficiencies uncovering the entire *Efa6* gene locus (*Efa6^Def^*), or different loss-of-function mutant alleles generated by genomic engineering (*Efa6^KO#1^*, *Efa6^GX6[w-]^*, *Efa6^GX6[w+]^*). In all these conditions, axon length at 6 HIV was increased compared to wild-type by at least 20% (Fig. 2D).

**Fig. 2.**
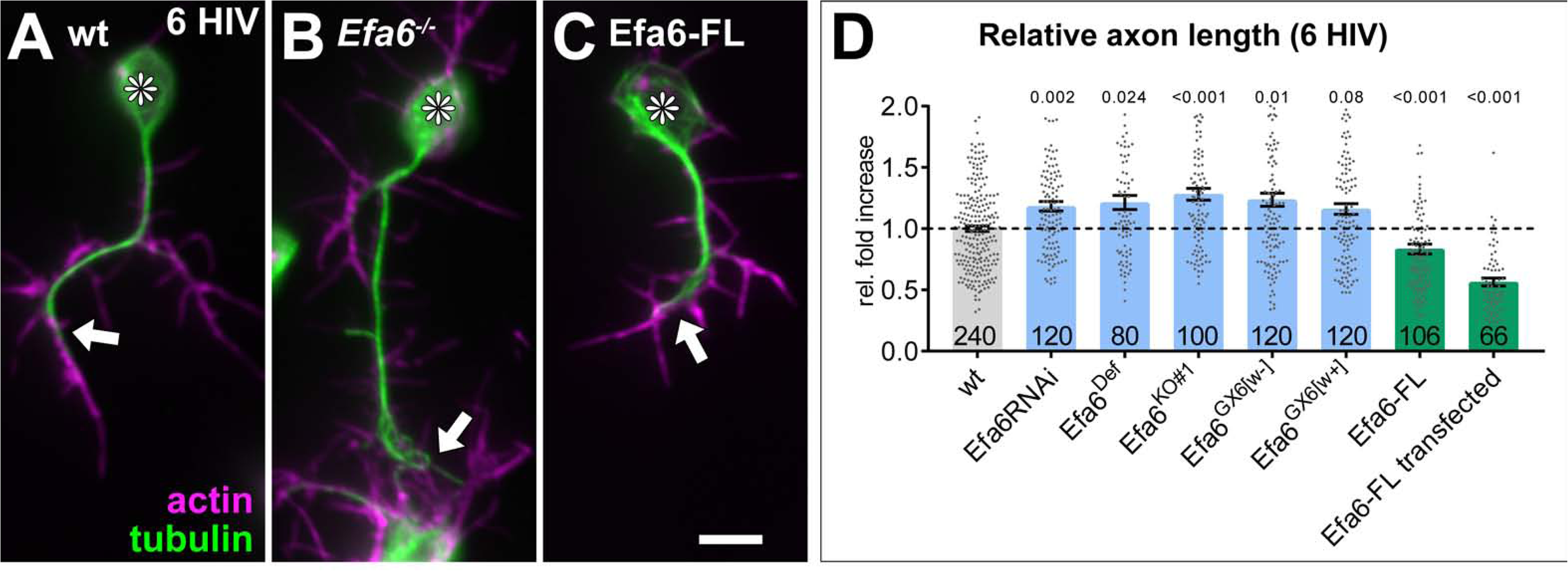
Efa6 regulates axonal length in primary *Drosophila* neurons. Examples of primary *Drosophila* neurons at 6HIV (A-C), all stained for actin (magenta) and tubulin (green); neurons are either wild-type controls (A), Efa6-deficient (B), expressing Efa6-FL::GFP (C); asterisks indicate cell bodies, arrows point at axon tips; the scale bar in C represents 10µm. Quantification of axon lengths at 6HIV (D); different genotypes are colour-coded: grey, wild-type controls; blue, different *Efa6* loss-of-function conditions; green, neurons over-expressing Efa6 variants; data represent fold-change relative to wild-type controls (indicated as horizontal dashed “ctrl” line); they are shown as single data points and a bar indicating mean ± SEM data; P values from Mann-Whitney tests are given above each column, sample numbers at the bottom of each bar. For raw data see T2, which can be downloaded here: <u>w.prokop.co.uk/Qu+al/RawData.zip</u>.

We then tested whether over-expression of Efa6 would cause the opposite effect, i.e. axon shortening or even loss. For this, we generated a transgenic *UAS-Efa6-FL-GFP* line and, in addition, developed methods to transfect *UAS*-constructs into *Drosophila* primary neurons (see Methods). When expressed pan-neuronally transgenic or transfected full-length *Efa6-FL::GFP* localised to cell bodies, axons and growth cones of primary neurons, as similarly observed with the endogenous protein (Figs.1B,C, S3B). The transgenic expression caused a ∼20% reduction in axon length, which was increased to ∼50% upon transfection (likely due to higher copy numbers of the expression construct; Fig. 2C, D). Furthermore, we observed an increase in the number of neurons without axons from ∼26% in *UAS-GFP*-transfected controls to ∼43% in Efa6-FL::GFP-positive neurons (Fig.3B).

**Fig. 3.**
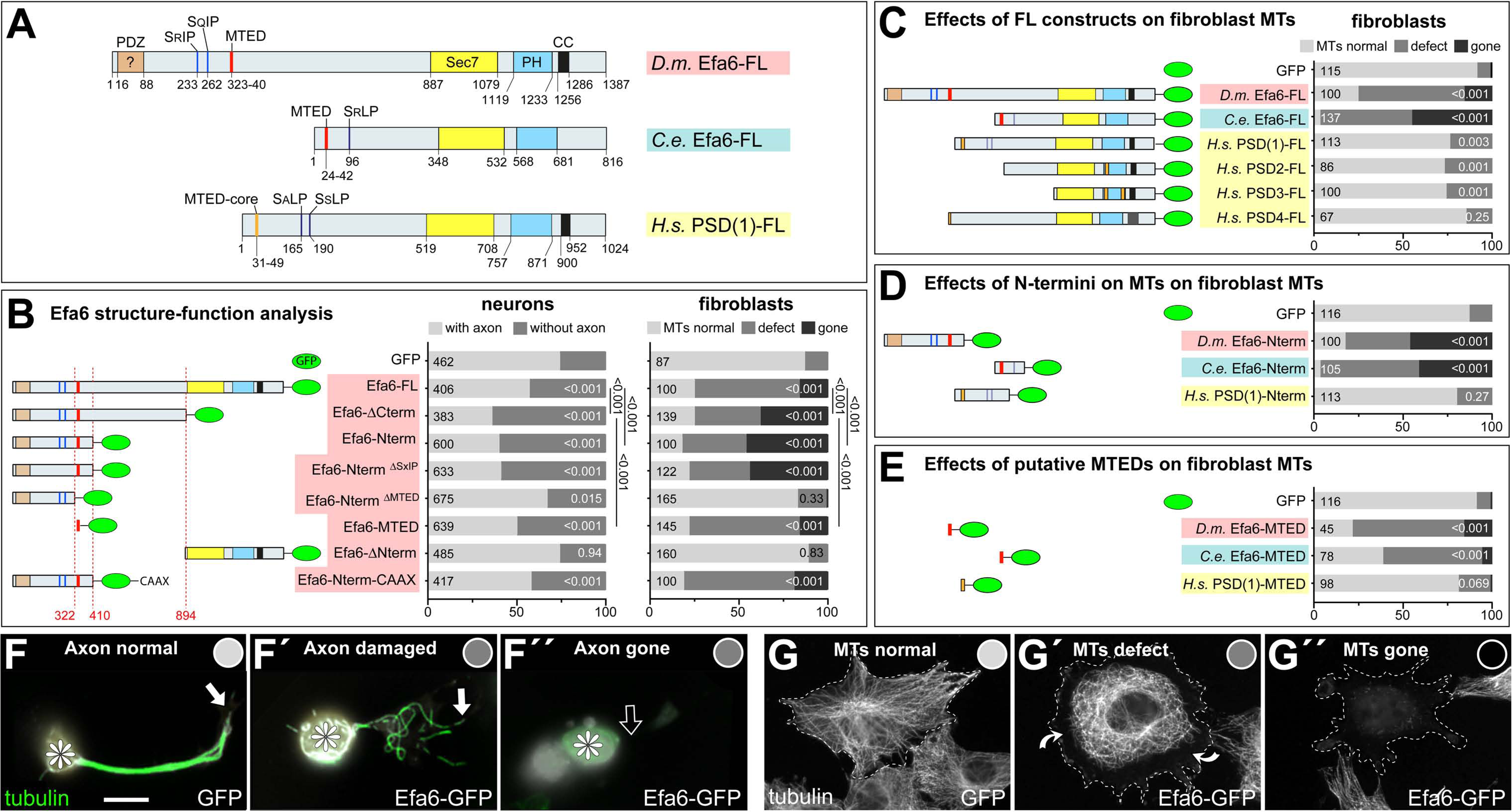
Efa6 domain and motif requirements for MT inhibition in neurons and fibroblasts. A) Schematics of *Drosophila melanogaster (Dm)* Efa6 (isoform C, CG31158), *Caenorhabditis elegans (Ce)* Efa6 (isoform Y55D9A.1a) and *Homo sapiens* (human) PSD1 (isoform 201/202, NP_002770.3), illustrating the positions (numbers indicate first and last residues) of the putative PSD95-Dlg1-ZO1 domain [PDZ; expected to anchor to transmembrane proteins (Ponting et al., 1997), but not mediating obvious membrane association in fibroblasts: Figs.S4C,D], SxIP/SxLP motifs (SRIP, SQIP, SALP, SSLP), the MT-binding domain (MTED), SEC7 domain, plekstrin homology (PH) domain and coiled-coil domain (CC). B) Schematics on the left follow the same colour code and show the *Dm*Efa6 constructs used in this study (dashed red lines indicate the last/first residue before/behind the truncation). Bar graphs on the right show the impact that transfection of these constructs had on axon loss in primary *Drosophila* neurons (dark grey in left graph) and on MT loss in fibroblasts (dark grey or black as indicated; for respective images see F and G below). Analogous fibroblast experiments as performed with *Drosophila* constructs were performed with full length constructs of *C. elegans* Efa6 and human PSDs (C), with N-terminal constructs (D) or synthetic MTEDs (E) of *Dm* and *Ce*Efa6 and of human PSD1. Throughout this figure, construct names are highlighted in red for *Drosophila*, light blue for *C. elegans* and yellow for *Homo sapiens*; all graph bars indicate percentages of neurons with/without axons (light/dark grey) and of fibroblasts with normal, reduced or absent MTs (light, medium, dark grey, respectively); numbers in the left end of each bar indicate sample numbers, on the right end the P values from Chi^2^ tests relative to GFP controls; numbers on the right of bars in B compare some constructs to Efa6-FL::GFP, as indicated by black lines. F-F’’) Primary neurons expressing Efa6-FL::GFP transgenically and stained for tubulin (asterisks, cell bodies; white arrows, axon tips; open arrow, absent axon). G-G’’) Fibroblasts expressing Efa6-FL::GFP and stained for tubulin; curved arrows indicate areas where MTs are retracted from the cell periphery; grey dots in F-G’’ indicate the phenotypic categories for each neuron and fibroblasts, as used for quantitative analyses in the graphs above. Scale bar at bottom of F refers to 10µm in F and 25µm in G. For raw data see T3, which can be downloaded here: <u>w.prokop.co.uk/Qu+al/RawData.zip</u>.

Together, these results suggest that Efa6 restricts axonal growth, comparable to reports for *C. elegans* Efa6 (*Ce*Efa6; Chen et al., 2015; Chen et al., 2011). The loss of whole axons upon Efa6-FL::GFP over-expression might suggest that Efa6 performs its morphogenetic roles by inhibiting MTs.

### Efa6 eliminates peripheral or even entire MT networks in mouse fibroblasts

To assess whether the negative impact of Efa6 on axon outgrowth might be through inhibiting MTs, we used NIH3T3 mouse fibroblasts as a heterologous cell system known to provide meaningful readouts for functional studies of *Drosophila* MT regulators (Alves-Silva et al., 2012; Beaven et al., 2015). When fibroblasts were analysed 24 hrs after transfection with *Efa6-FL-GFP*, we found a graded depletion of MT networks depending on Efa6-FL::GFP protein levels (shown and quantified in Fig.S5). At moderate expression levels, Efa6-FL::GFP localised along the circumference and in areas of membrane folds (open arrow heads in Figs.S5B), and MTs tended to be lost predominantly from the cell fringes (curved arrows in Figs.S5B and S7B). At high expression levels, Efa6-FL::GFP became detectable in the cytoplasm and even nucleus (double-chevrons in Fig.S5C), suggesting that membrane-association might become saturated. In these cases, MTs were completely absent (Fig.S5C). When quantifying these MT phenotypes across all transfected fibroblasts, there was a strong increase in MT network defects and depletion upon Efa6-FL::GFP expression as compared to GFP controls (Fig. 3B).

When performing live analyses, we consistently observed that growing MTs labelled with EB3::mCherry extended to the very cell fringes of control fibroblasts (Suppl. Movie M1), whereas MTs in fibroblasts transfected with Efa6-FL::GFP showed a very different behaviour: hardly any MTs polymerised into areas along the rim where Efa6 was enriched but stopped at the border, often accompanied by Efa6-FL::GFP accumulation at MT plus ends at the invasion site (Suppl. Movie M2).

Taken together, also these data suggest that Efa6 inhibits MTs. The fibroblast experiments suggest that Efa6 is membrane-associated and excludes MTs from this position, and the studies in fly neurons indicate the relevance of such functions for neuronal morphogenesis. The combined use of mouse fibroblasts and *Drosophila* primary neurons provides therefore a robust system with informative readouts for MT loss - ideal to carry out a systematic structure-function analysis of Efa6.

### The N-terminal 18aa motif of Efa6 is essential for microtubule-inhibiting activity of Efa6

A detailed analysis of the domain structures of Efa6 proteins from 30 species revealed that C-termini of almost all species contain a putative pleckstrin homology domain (PH; potentially membrane-associating; Macia et al., 2008), a Sec7 domain (potentially activating Arf GTPases; D’Souza-Schorey and Chavrier, 2006; Huang et al., 2009) and a coiled-coil (CC) domain (Franco et al., 1999; Figs.3A, S2). In contrast, the N-termini are mainly unstructured and reveal enormous length differences among species. Accordingly, phylogenetic relationship analyses comparing either full-length or N-terminal Efa6, show that chordate proteins are rather distant from invertebrates, and that arthropods form a clear subgroup within the invertebrates (Fig. S1A,B). None of the identifiable N-terminal domains/motifs is particularly well conserved (details in Fig.S2). For example, the *Drosophila* N-terminus contains (1) a putative PDZ domain (aa16-88; mainly found in insect versions of Efa6), (2) two SxIP motifs (aa 233-6 and 262-5; found primarily in Efa6 of flies, some other insects and molluscs; some vertebrate/mammalian species display derived SxLP motifs), and (3) a motif of 18aa displaying 89% similarity with a motif in the N-terminus of *Ce*Efa6 suggested to be involved in MT inhibition (O’Rourke et al., 2010; conserved in nematodes, arthropods and molluscs).

To assess potential roles of the *Drosophila* 18aa motif (from now on referred to as MT elimination domain, MTED), we generated a series of GFP-tagged N-terminal constructs (Fig.3B): *Efa6-ΔCterm-GFP* (encoding the entire N-terminal half upstream of the Sec7 domain), *Efa6-Nterm-GFP* (restricting to the N-terminal part containing all the identified functional domains), *Efa6-Nterm^ΔMTED^-GFP* (lacking the MTED) and *Efa6-MTED-GFP* (encoding only the MTED). All these N-terminal Efa6 variants showed the same localisation pattern throughout neurons (Fig.S3C,D,F,G), and in the cytoplasm and nucleus of fibroblasts (Fig.S4C,D,F,G). Cytoplasmic and nuclear localisations occurred even at low expression levels, indicating that the absent C-terminus (and likely PH domain within) usually mediates membrane association. This nuclear localisation occurs in the absence of any predicted N-terminal nuclear localisation sequences (Figs.3A, S2A), likely reflecting a known artefact of GFP-tagged proteins (Alves-Silva et al., 2012; Seibel et al., 2007).

In spite of their very similar localisation patterns, the functional impact of these constructs was clearly MTED-dependent: only constructs containing the MTED (Efa6-ΔCterm::GFP, Efa6-Nterm::GFP and Efa6-MTED::GFP) caused strong axon loss in neurons and MT network depletion in fibroblasts, whereas Efa6-Nterm^ΔMTED^::GFP behaved like GFP controls (Figs.3B; S3C,D,F,G; S7C,D,G,H).

In addition, we assessed potential roles of the two SxIP sites predicted to bind EB proteins (Honnappa et al., 2009; Fig.3A). Accordingly, we found that Efa6-Nterm::GFP tip-tracks and that Efa6-FL::GFP accumulates at sites where EB3-enriched MTs get in contact (Suppl. Mov. M2 and 3). Such binding to EBs at MT plus ends, might enhance Efa6’s ability to capture MTs for inhibition. However, when replacing each of the two SxIP motifs by four alanines (Efa6-Nterm^ΔSxIP^::GFP), the construct still induced a strong axon loss and MT network depletion in fibroblasts (Figs.3B, S3E, S4E, S7F). Similar observations were reported for Kif2C, which clearly tip-tracks MTs through binding EB1, but does not require this property for its MT depolymerising activity (Moore et al., 2005).

Taken together, our results pinpoint the MTED as the key mediator of MT-depleting functions of *Drosophila* Efa6, suggesting this function to be conserved between flies and *C. elegans*.

### The MTED is a good predictor of MT-inhibiting function directly affecting MT polymerisation

To assess whether the MTED motif is a good predictor for MT-inhibiting capabilities of Efa6 family members, we used 12 different constructs comprising: full length versions of (1) *Ce*Efa6, (2) *Drosophila* Efa6 and (3-6) all four human PSDs (Fig. 3C), as well as N-terminal versions of (7) *Ce*Efa6, (8) fly Efa6 and (9) human PSD1 (Fig. 3D). Furthermore, we deduced a MTED consensus sequence from 39 *Efa6* genes (details in Fig. S2C), identified the most likely human MTED-like sequence (position 31-49aa of PSD1; MTED-core in Fig. 3A) and synthesised codon-optimised versions of (10) this human as well as the (11) fly and (12) worm MTEDs (Figs. S2B,C). When transfected into fibroblasts, we found that all 6 fly/worm constructs had strong MT-inhibiting properties, whereas none of the 6 human constructs (PSD1-4 full length, PSD1-Nterm, PSD1-MTED-like) showed MT collapse (Fig.3C-E). Therefore, the presence of a well conserved canonical MTED seems to be a good predictor for MT-inhibiting capabilities of Efa6 proteins.

To gain insights into the mechanisms through which MTEDs might act, we carried out a series of *in vitro* experiments. Purified Efa6-Nterm::GFP clearly associated with MTs *in vitro* (Fig. S8A), but failed to reconstitute any MT-inhibiting activity (Fig. S8B). We therefore tested the same protein in *Xenopus* oocyte extract to assess potential co-factor requirements, but saw again no activity (Fig. S8C) - in spite of the fact that injection of a corresponding mRNA into *Xenopus* oocytes caused strong cell division phenotypes (Fig. S8D,E). We suspected problems with recombinant expression of the protein, which is predicted to have large disordered regions, and instead used synthetic MTED peptide. We found that MTED-coated sepharose beads pulled down unpolymerised tubulin (Fig.4B), indicating a direct interaction of the peptide with tubulin. Furthermore, addition of synthetic MTED peptide to a MT growth assay, resulted in strong suppression of MT polymerisation in a dose-dependent manner (Fig.4A).

**Fig. 4.**
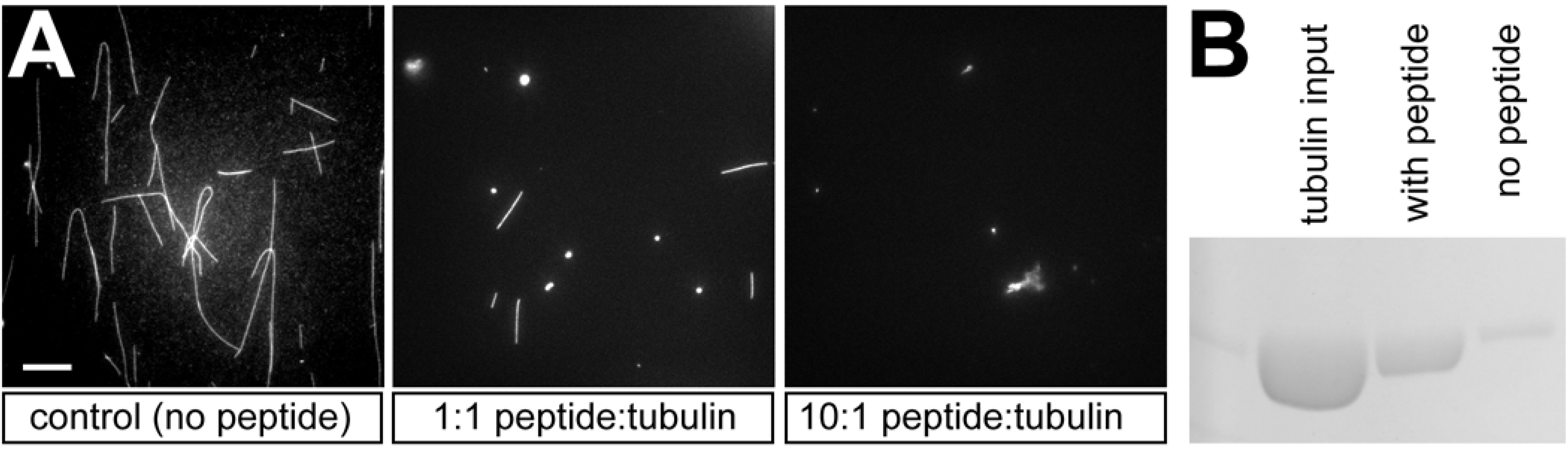
EFA6 peptide interacts directly with α/β-tubulin and inhibits microtubule growth. A) End point of microtubule growth assay in the presence of no MTED peptide, or a 1:1 and 10:1 molar ratio of MTED peptide to tubulin. Scale bar in A refers to 10 µm. B) Pull-down of tubulin by sepharose beads coated with MTED peptide via cyanogen bromide coupling and control beads with no MTED peptide attached.

Taken together, the MTED exists primarily in Efa6 homologues of invertebrate species and its presence correlates with MT-inhibiting properties of these proteins. This conclusion is strongly supported by our finding that *Drosophila* MTED directly interferes with MT polymerisation, which can explain why MTs fail to enter Efa6-enriched areas in fibroblasts (Suppl. Mov. M2).

### The C-terminal domain restricts the microtubule-inhibiting activity of Efa6 to the cortex

Our structure-function analyses strongly suggested that Efa6 is membrane-associated. This is further supported by a membrane ruffle phenotype we observed in fibroblasts when expressing Efa6-FL::GFP or the C-terminal derivative Efa6-ΔNterm::GFP (Figs.3B; curved open arrows in Figs.S4B,H and S6B,D). Efa6-ΔNterm::GFP had no obvious effects on MT networks (Fig. S7I), and its membrane ruffling phenotype likely reflects an evolutionarily conserved function of the Efa6 C-terminus through its Sec7, PH and/or CC domains (Derrien et al., 2002; Franco et al., 1999; Macia et al., 2008). Accordingly, we find the same membrane ruffling when expressing PSD1-FL::GFP (curved open arrows in Fig. S6E).

However, even if the C-terminus plays no active role in the MT inhibition process, it still regulates this function. This is suggested by the Efa6-ΔCterm::GFP and Efa6-Nterm::GFP variants which lack the C-terminus (Fig.3B), fail to associate with the cortex (Fig.S4C,D,F,G), do not cause ruffling (Fig.S6C), but induce MT phenotypes far stronger than Efa6-FL::GFP does (Figs.3B; S7C,D,H *vs* B). To assess whether lack of membrane tethering could explain this phenotypic difference, we generated the Efa6-Nterm::GFP::CAAX variant (Fig.3B) where Efa6-Nterm::GFP is fused to the membrane-associating CAAX domain (Hancock et al., 1991); this addition of CAAX changed the properties of Efa6-Nterm::GFP back to Efa6-FL::GFP-like behaviours (Figs.3B; S4B,I; S7B,E): the hybrid protein localised to the cortex in fibroblasts and had only a moderate MT phenotype, and also the axon loss phenotype was mild. Also in live analyses, the CAAX construct reproduced the effect of excluding MTs from Efa6-N-term::GFP::CAAX-enriched areas (Suppl. Mov. M4). These findings confirm membrane tethering as an important regulatory feature restricting Efa6 function.

Taken together, our structure-function data clearly establish *Drosophila* Efa6 as a cortical collapse factor: its N-terminal MTED blocks polymerisation which of MTs at the cortex via the Efa6 C-terminus.

### Efa6 negatively regulates MT polymerisation at the growth cone membrane and in filopodia

We next asked how Efa6’s cortical collapse function relates to the observed axon growth phenotypes. For this, we focussed on growth cones (GCs) as the sites where axons extend; this extension requires the splaying of MTs from the axonal bundle tip at the base of GCs to explore the actin-rich periphery (Dent et al., 2011; Lowery and van Vactor, 2009; Prokop et al., 2013).

In GCs of primary neurons at 6 HIV, loss of Efa6 caused an increase in MT polymerisation events: the total number of Eb1 comets was increased as compared to wild-type controls (Fig. 5I). Eb1::GFP comets in *Efa6* mutant neurons frequently persisted when reaching the GC periphery, where they could occasionally be observed to undergo curved extensions along the periphery (Suppl. Mov. M7and SM8). Comet velocity was unaffected (∼0.3 μm/s), but the lifetime of Eb1::GFP comets was ∼1.4 times longer in mutant GCs, with the dwell time of Eb1 comets at the tip of filopodia being increased from 2.10s +/- 0.24 s in wild-type to 6.26s +/-0.40s in *Efa6* mutant neurons (Fig. 5M); we even observed cases where comets at the tips of filopodia were moving backwards, seemingly pushed back by the retracting filopodial tip (Suppl. Mov. M7 and M8). In agreement with the increased lifetime, more microtubules invaded growth cone filopodia in *Efa6* mutant neurons, as quantified by counting filopodia that contained EB1comets or MTs (Fig.5D,E,J,K; note that the total number of filopodia per GC was in the range of 10-11 for both wild-type and *Efa6*; not shown). Transgenically expressed Efa6-FL::GFP caused the opposite effect, i.e. a reduction in the number of GC filopodia containing Eb1 comets or MTs (Fig. 5G-L; green columns).

**Fig. 5.**
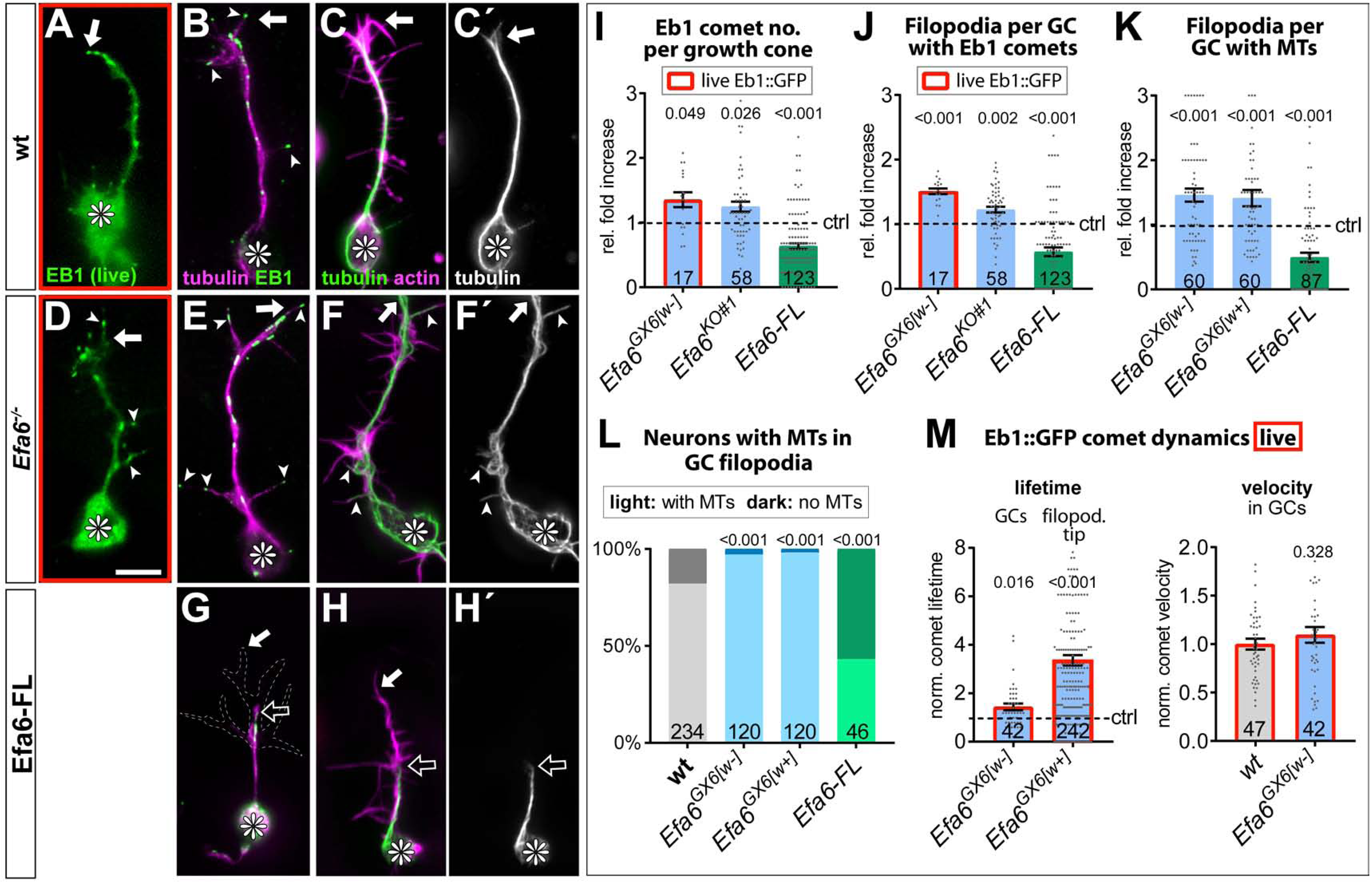
Efa6 regulates MT behaviours in GCs. A-H’) Examples of primary neurons at 6HIV which are either wild-type controls (top), Efa6-deficient (middle) or expressing Efa6-FL::GFP (bottom); neurons were either imaged live for Eb1::GFP (green in A,D) or fixed and labelled for Eb1 and tubulin (B,E,G; as colour-coded) or actin and tubulin (C,F,H; as colour coded; tubulin shown as single channel image on the right); asterisks indicate cell bodies, white arrows the tips of GCs, open arrows the tips of MT bundles and arrow heads MTs or Eb1 comets in filopodial processes; the GC in G is outlined with a white dashed line; scale bar in D represents 5µm in all images. I-M) Quantitative analyses of MT behaviours in GCs of such neurons as indicated above each graph. Different genotypes are colour-coded: grey, wild-type controls; blue, different *Efa6* loss-of-function conditions; green, neurons over-expressing Efa-FL. The graph in L shows percentages of neurons without any MTs in shaft filopodia (dark shade) versus neurons with MTs in at least one filopodium (light shade; P values above bars assessed via Chi^2^ tests), whereas all other graphs show single data points and a bar indicating mean ± SEM, all representing fold-increase relative to wild-type controls (indicated as horizontal dashed “ctrl” line; P values above columns from Mann-Whitney tests. The control values in M (dashed line) equate to an Eb1 comet life-time of 2.10s ± 0.24SEM in filpododia and 5.04s ± 0.60SEM in growth cones, and a comet velocity of 0.136 μm/s ± 0.01SEM.Throughout the figure, sample numbers are shown at the bottom of each bar and data obtained from live analyses with Eb1::GFP are framed in red. For raw data see T4, which can be downloaded here: <u>w.prokop.co.uk/Qu+al/RawData.zip</u>.

Next, we investigated MT dynamics in axon shafts. In contrast to GCs, MTs in the axon shaft are organised into bundles, hence kept away from the membrane. Accordingly, neither loss-nor gain-of-function had an obvious effect on Eb1 comet numbers, lifetimes, velocities and directionalities (Fig. 6A-D). However, like in GCs, there was a strong increase in filopodia along the shaft that contained MTs when Efa6 was absent, and a strong decrease when over-expressing Efa6-FL::GFP (Fig. 6E-G).

**Fig. 6.**
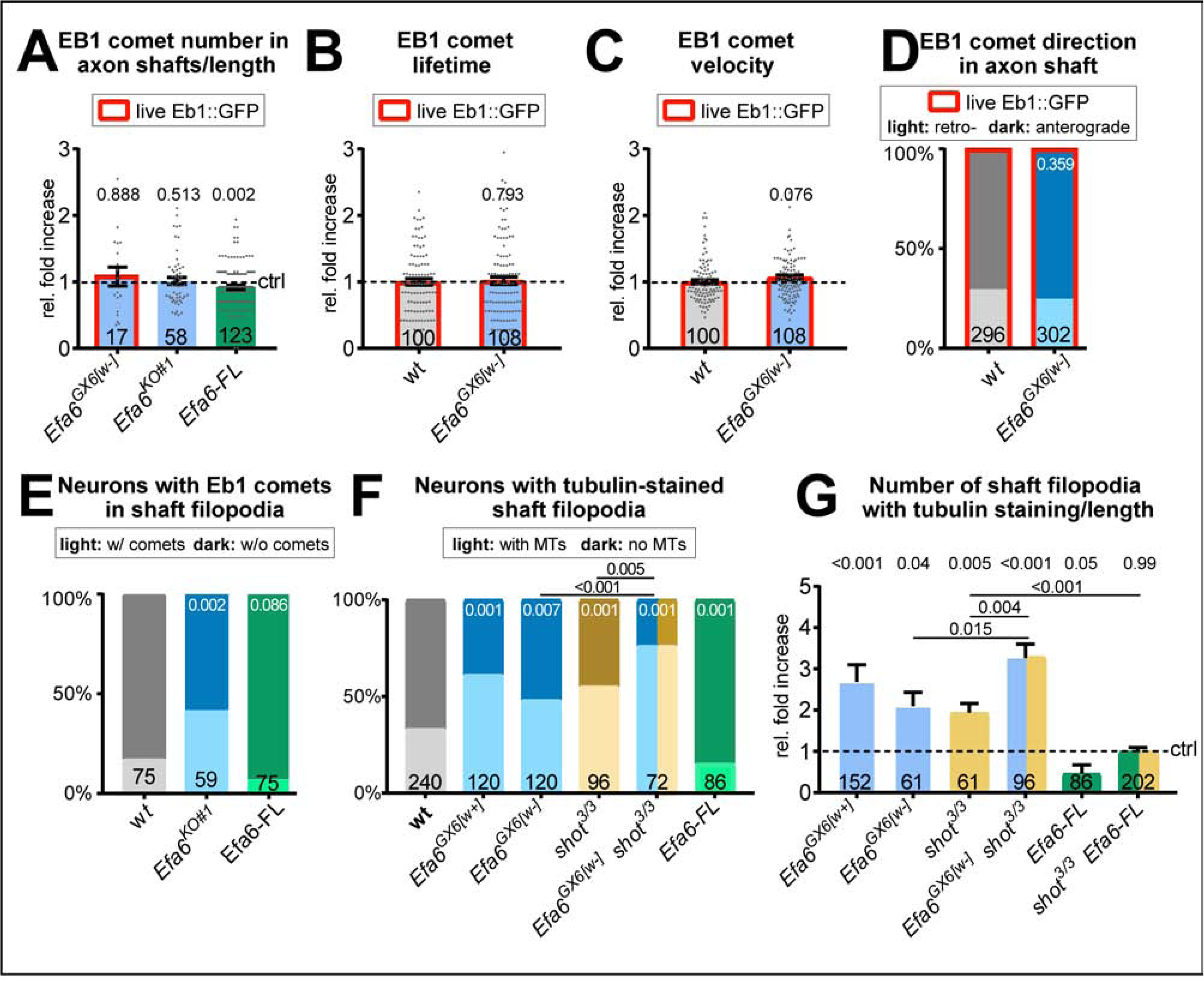
Loss of Efa6 promotes MT entry into axon shaft filopodia. Quantitative analyses of MT behaviours in axon shafts, as indicated above each graph; bars are colour-coded: grey, controls; blue, different *Efa6* mutant alleles; green, neurons over-expressing Efa-FL::GFP or Efa6::CAAX::GFP; orange, *shot* mutant allele; red outlines indicate live imaging data, all others were obtained from fixed specimens; numbers at the bottom of bars indicate sample numbers, above bars P values. A-C,G) Fold-changes relative to wild-type controls (indicated as horizontal dashed “ctrl” line) shown as single data points and a bar indicating mean ± SEM; P values were obtained via Mann-Whitney tests; control values (dashed line) in B and C equate to an Eb1 comet lifetime of 7.18 s ± 0.35 SEM and a velocity of 0.169 μm/s ± 0.01SEM. D-F) Binary parameters (light *versus* dark shades as indicated) provided as percentages; P values were obtained via Chi^2^ tests. For raw data see T5, which can be downloaded here: <u>w.prokop.co.uk/Qu+al/RawData.zip</u>.

Taken together, our data are consistent with a model in which Efa6 primarily inhibits explorative MTs that leave the axon bundle in either GCs or axon shafts and polymerise towards the cell membrane or into filopodia. Surplus MTs in the periphery of GCs can explain the extra axonal growth we observed (Fig.2).

### Efa6 negatively influences axon branching

We hypothesised that an increase in explorative MTs could also cause a rise in axon branching (see Introduction), either by inducing GC splitting through parallel growth events in the same GC (Acebes and Ferrus, 2000), or by seeding new collateral branches along the axon shaft (Kalil and Dent, 2014; Lewis et al., 2013). To test this possibility, we studied mature primary neurons at 5 days *in vitro* (DIV). We found that *Efa6^KO#1^* homozygous mutant neurons showed almost double the number of collateral branches as observed in wild-type neurons, whereas expression of Efa6-FL::GFP reduced branching by 21% (Fig.7A-C,E). This reduction is mediated by the Efa6 N-terminus, since expression of Efa6-Nterm::GFP::CAAX caused a similar degree in branch reduction (Fig.7D,E).

**Fig. 7.**
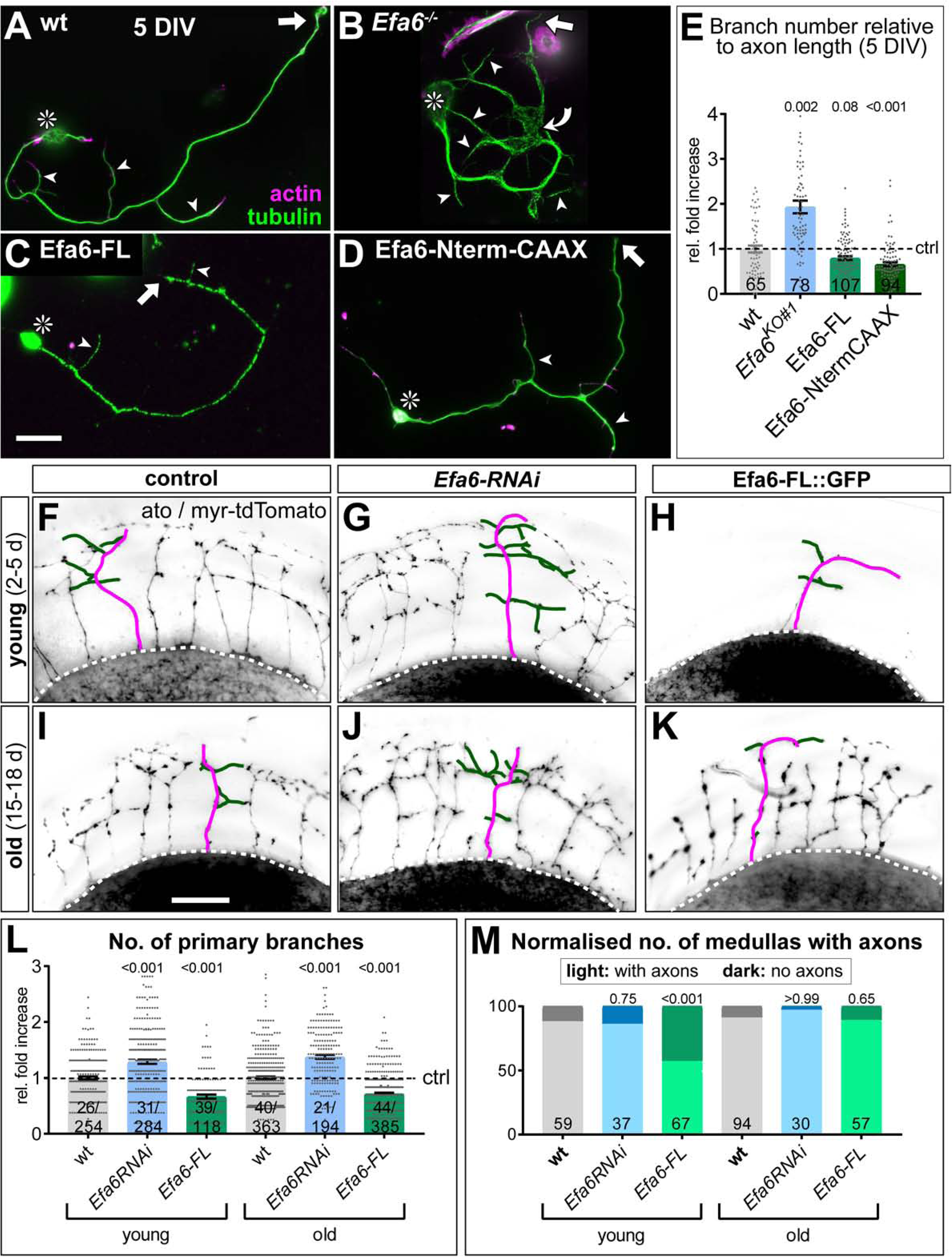
Efa6 regulates axon branching in in primary *Drosophila* neurons and adult fly brains. Examples of primary *Drosophila* neurons at 5DIV (A-D), all stained for actin (magenta) and tubulin (green); neurons are either wild-type controls (A), Efa6-deficient (B), expressing Efa6-FL::GFP (C), or expressing Efa6-Nterm-CAAX::GFP (D); asterisks indicate cell bodies, arrows point at axon tips, arrow heads at axon branches, the curved arrow at an area of MT disorganisation; the scale bar in C represents 20µm. Quantification of axonal branch numbers (E); different genotypes are colour-coded: grey, wild-type controls; blue, different *Efa6* loss-of-function conditions; green, neurons over-expressing Efa6 variants; data represent fold-change relative to wild-type controls (indicated as horizontal dashed “ctrl” line); they are shown as single data points and a bar indicating mean ± SEM data; P values from Mann-Whitney tests are given above each column, sample numbers at the bottom of each bar. F-K) Brains (medulla region of the optic lobe in oblique view) of young (2-5 d after eclosure; top) and old flies (15-18 d; bottom) driving *UAS-myr-tdTomato* via the *ato-Gal4* driver in dorsal cluster neurons (example neurons are traced in magenta for axons and green for side branches); flies are either wild-type controls (left) or display *ato-Gal4*-driven knock-down of Efa6 (middle) or over-expression of Efa6-FL::GFP (right). L,M) Quantification of data for wild-type (wt; grey), Efa6 knock-down (blue) and Efa6-FL::GFP over-expression (green): L) shows the number of primary branches per axon as fold-change relative to wild-type controls (indicated as horizontal dashed “ctrl” line; data are shown as bars indicating mean ± SEM accompanied by single data points, all normalised to wt); M) displays the number of medullas with axons (light colours) or without axons (dark colours) shown as a percentages in young and old flies. In all graphs, sample numbers at the bottom (in L number of medullas before and number of axons after slash) and P values from Mann-Whitney (G) or Chi^2^ tests (H) above each column. Scale bar in D represents 60µm in all figures. For raw data see T6, which can be downloaded here: <u>w.prokop.co.uk/Qu+al/RawData.zip</u>.

To extend these studies to neurons *in vivo*, we studied dorsal cluster neurons, a subset of neurons with stereotypic axonal projections in the optic lobe of adult brains (Fig.7F-K, see Methods; Hassan et al., 2000; Voelzmann et al., 2016b). To manipulate Efa6 levels in these neurons, either *Efa6-RNAi* or *Efa6-FL-GFP* was co-expressed with the membrane marker *myr-tdTomato*. Their axon branches were assessed in brains of young and old flies (2-5 d and 15-18 d after eclosure from the pupal case, respectively; Figs.7F-K). We found that *Efa6* knock-down in dorsal cluster neurons caused a significant increase in branch numbers by 29% in young and by 38% in old brains, whereas over-expression of Efa6::GFP strongly decreased branch numbers by 33% in young and 28% in old brains, respectively (Fig.7L).

In these experiments, Efa6-FL::GFP expression had an intriguing further effect: Only 57% of young brains had any axons in the medulla region, compared to 88% in controls (Figs.7H,M, S9B). However, in the older Efa6-FL::GFP expressing fly brains, the axons were eventually present (Fig.7K,M, S9D). We concluded that this phenotype reflected delayed outgrowth, which is also consistent with the decrease in axon growth observed upon Efa6 over-expression in primary neurons (green bars in Fig.2D).

Taken together, our data indicate a physiologically relevant role of Efa6 as negative regulator of axonal branching, mediated through its N-terminus, most likely via its function as cortical collapse factor.

### *Efa6* maintains axonal MT bundle integrity in cultured neurons

Apart from changes in growth and branching, we noticed that a significant amount of Efa6-depleted neurons displayed axons with swellings where MTs lost their bundled conformation and were arranged into intertwined, criss-crossing curls instead (Fig.S10; arrowheads in Fig. 8D-F). To quantify the strength of this phenotype, we measured the area of MT disorganisation relative to axon length (referred to as ‘MT disorganisation index’, MDI; Qu et al., 2017). MDI measurements in *Efa6* mutant neurons revealed a mild 1.3 fold increase in MT disorganisation in young neurons which gradually worsened to 2.3 fold at 5 DIV and ∼4 fold at 10 DIV (Fig.8A-F,I; all normalised to controls).

**Fig. 8.**
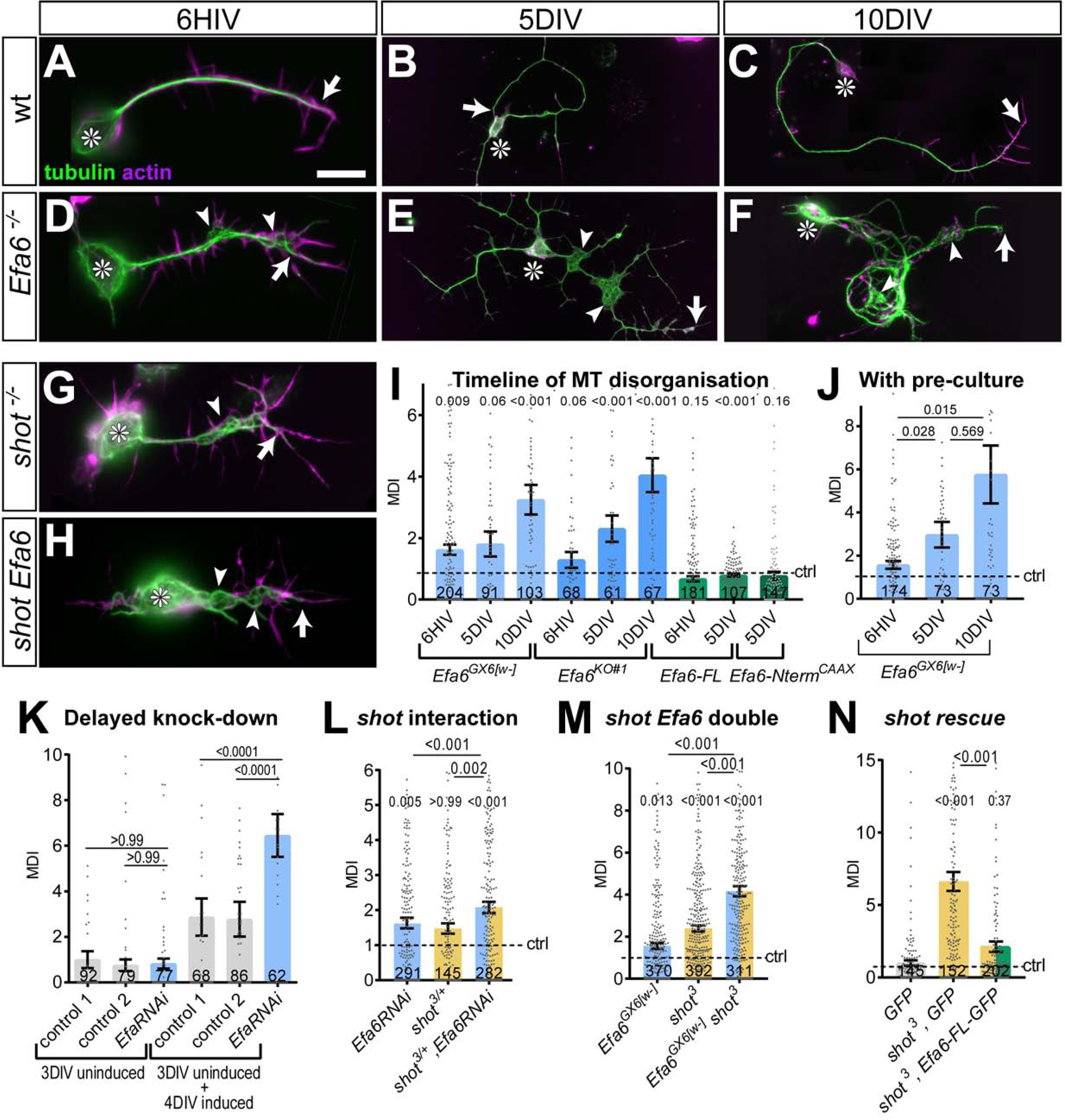
Efa6 helps to maintain axonal MT bundles in *Drosophila* neurons. A-H) Images of primary neurons at 6HIV (left), 5DIV (middle) and 10DIV (right), stained for tubulin (green) and actin (magenta), derived from embryos that were either wild-type (wt, A-C), *Efa6* null mutant (D-F), homozygous for *shot^3^*(G) or *shot^3^ Efa6^GX6[w-]^* double-mutant (shot Efa6, H); arrows point at axon tip, arrow heads at areas of MT disorganisation, and asterisks indicate the cell bodies; the scale bar in A represents 10µm for 6HIV neurons and 25µm for 5DIV and 10DIV neurons. I-N) Quantitative analyses of MT disorganisation (measured as MT disorganisation index, MDI) in different experimental contexts (as indicated above graphs); different genotypes are colour-coded: grey, wild-type controls; blue, *Efa6* loss-of-function; light/dark orange, *shot^3^*in hetero-/homozygosis; green, neurons over-expressing Efa-FL::GFP or Efa6-NtermCAAX::GFP; data are shown as single data points and a bar indicating mean ± SEM, all representing fold-change relative to wild-type controls (indicated as horizontal dashed “ctrl” line); P values from Mann-Whitney tests in I, J) and Kruskal–Wallis one-way ANOVA with *post hoc* Dunn’s test in K-N) are given above each column, sample numbers at the bottom of each bar. Control 1 in (K) is *tub-Gal80*, *elav-Gal4* alone and control 2 *UAS-Efa6RNAi*. For raw data see T7, which can be downloaded here: <u>w.prokop.co.uk/Qu+al/RawData.zip</u>.

The observed gradual increase in phenotype could be the result of a genuine function of *Efa6* not only during axon growth but also their subsequent maintenance. Alternatively, it could be caused by maternal gene product deposited in the mutant embryos by their heterozygous mothers (Prokop, 2013b); such maternal *Efa6* could mask mutant phenotypes at early stages so that they become apparent only after most Efa6 has degraded. To assess the latter possibility, we used a pre-culture strategy to remove potential maternal Efa6 (see Methods; Prokop et al., 2012; Sánchez-Soriano et al., 2010). When plating neurons after 5 days of pre-culture, we still found a low amount of MT disorganisation in young neurons and a subsequent gradual increase to severe phenotypes over the following days (Fig.8J).

This finding argues for a continued role of Efa6 in preventing MT disorganisation during development as well as in mature neurons. To further test this possibility, we used a temperature-based conditional knock-down technique (*elav-GAL4 UAS-Efa6-RNAi UAS-Gal80^ts^*abbreviated to *elav/Efa6^IR^/Gal80^ts^*; see Methods): the *elav/Efa6^IR^/Gal80^ts^* neurons were grown without knock-down (19°C) for 3 days, a stage at which they have long undergone synaptic differentiation (Küppers-Munther et al., 2004; Prokop et al., 2012); at that point, we found no difference in MT disorganisation between non-induced construct-bearing cells and control neurons (Fig. 8K). After this period, cells were grown for another four days under knock-down conditions (27°C), and then fixed on day seven. At this point, MT disorganisation in the *elav/Efa6^IR^/Gal80^ts^* neurons was significantly increased over control neurons (Fig. 8K), indicating that Efa6 is not only required during development but also during later maintenance to prevent MT disorganisation.

In contrast to increased MT disorganisation upon functional loss of Efa6, expression of Efa6-FL::GFP or Efa6-Nterm::GFP::CAAX showed a tendency to reduce MT disorganisation even below the baseline levels measured in control cells (cultured in parallel without the expression construct; Fig.8I), arguing that also this role of Efa6 is likely due to the cortical collapse function of Efa6 (see Discussion).

### *Efa6* maintains axonal MT bundle integrity *in vivo*

We then assessed whether a role of Efa6 in MT bundle maintenance is relevant *in vivo*. For this, we studied a subset of lamina neurons, which project prominent axons in the medulla of the adult optic lobe (Prokop and Meinertzhagen, 2006). We labelled MTs in these axons by expressing *α-tubulin84B-GFP* either alone (*GMR-tub* controls), or together with *Efa6^RNAi^* to knock down *Efa6* specifically in these neurons (*GMR-tub-Efa6^IR^*; see Methods for details).

When analysing aged flies at 26-27 days, we found that *Efa6* knock-down caused a doubling in the occurrence of axonal swellings with disorganised axonal MTs: the average of total swellings per column section was increased from 0.3 in controls to 0.65 swellings upon Efa6 knock-down; about a third of these contained disorganised MTs (*GMR-tub-Efa6^IR^*: 0.23 per column section; *GMR-tub*: 0.13; Fig.9). These data demonstrated that our findings in cultured neurons are relevant *in vivo*.

**Fig. 9.**
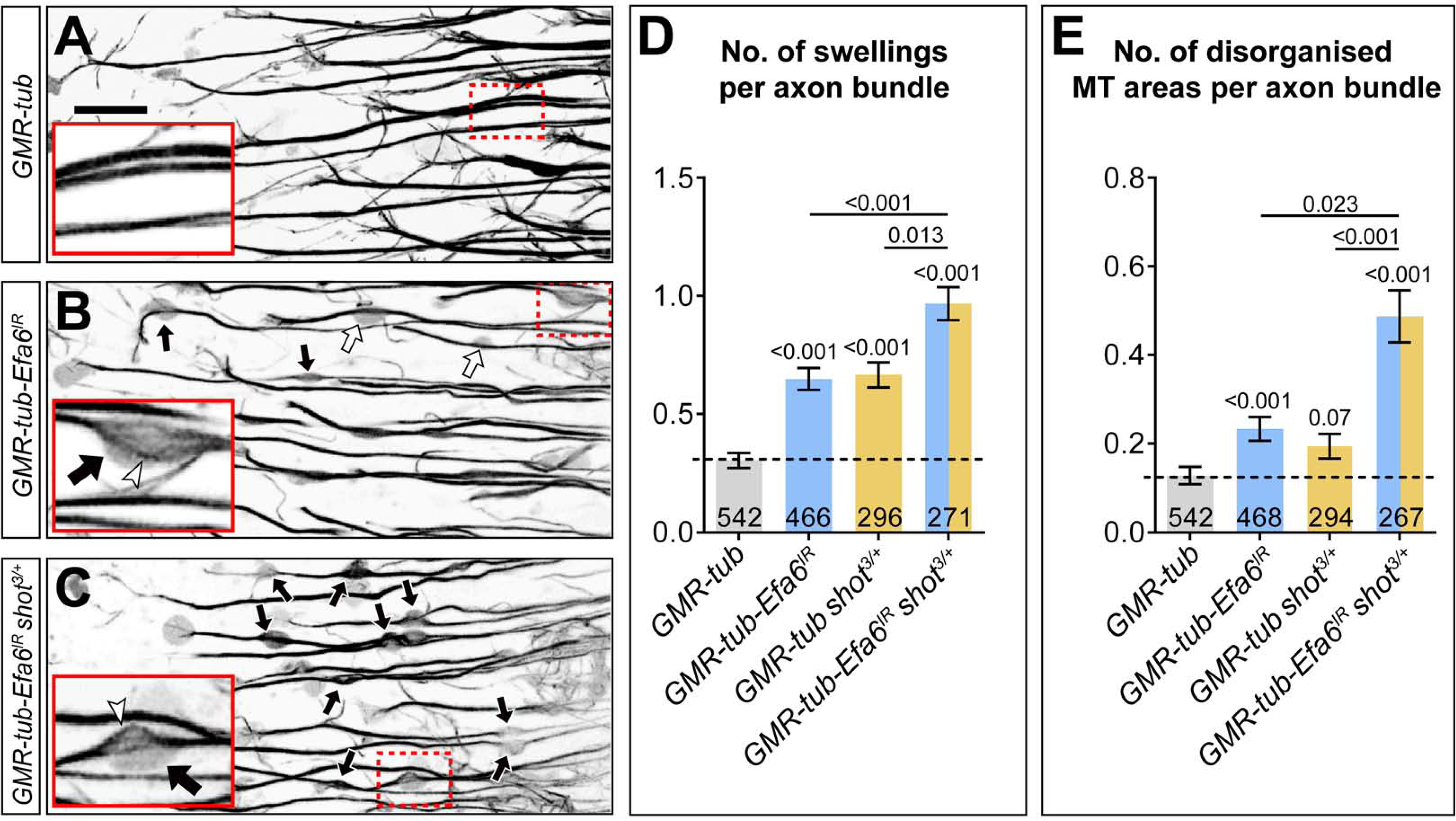
Efa6 is required for axonal MT bundle maintenance in adult fly brains. A-C) Medulla region of adult brains at 26-27 days after eclosure, all carrying the *GMR31F10-Gal4* driver and *UAS-α-tubulin84B-GFP* (*GMR-tub*) which together stain MTs in a subset of lamina neuron axons that terminate in the medulla. The other specimens co-express *Efa6-RNAi* either alone (*GMR-tub-Efa6^IR^*in B) or combined with a *shot^3/+^* heterozygous mutant background (*GMR-tub-Efa6^IR^ shot^3/+^* in C). White/black arrows indicate axonal swellings without/with MT disorganisation; rectangles outlined by red dashed lines are shown as 2.5 fold magnified insets where white arrow heads point at disorganised MTs; the scale bar in A represents 15µm in all images. D, E) Quantitative analyses of all axonal swelling (D) or swellings with MT disorganisation (E); different genotypes are colour-coded as indicated; bars show mean ± SEM, all representing fold-change relative to wild-type controls (indicated as horizontal dashed line); P values from Kruskal–Wallis one-way tests are given above each column, sample numbers at the bottom of each bar. For raw data see table T8, which can be downloaded here: <u>w.prokop.co.uk/Qu+al/RawData.zip</u>.

We propose therefore that Efa6 provides a quality control mechanism that prevents MT disorganisation by inhibiting only MTs that have escaped axonal bundles. This model would also be consistent with the slow onset and gradual increase of MT disorganisation we observed upon Efa6 deficiency (Fig.8I,J).

### Efa6 and Shot promote MT bundles through complementary mechanisms

If Efa6 provides a quality control mechanism that “cleans up” explorative MTs, it should act complementary to other factors that “prevent” explorative MTs by actively keeping them in axonal bundles. Very powerful preventive factors in both mammals and fly are the spectraplakins (Bernier and Kothary, 1998; Dalpe et al., 1998; Voelzmann et al., 2017). In *Drosophila*, spectraplakins are represented by the single *short stop* (*shot*) gene; *shot* deficiency causes a severe increase in axonal off-track MTs and MT disorganisation (Alves-Silva et al., 2012; Qu et al., 2017; Sánchez-Soriano et al., 2009).

To study potential mutual enhancement of *Efa6* and *shot* mutant phenotypes, we first determined numbers of MTs in axonal shaft filopodia: both single-mutant conditions showed a strong enhancement of filopodial MTs (blue *vs.* orange bars in Fig.6F,G); this phenotype was substantially further increased in *shot^3^ Efa6^GX6[w-]^* double-mutant neurons (orange/blue bars in Fig. 6F,G). *Vice versa*, when transfecting *Efa6-FL-GFP* to boost the hypothesised “cleaning-up” function, the *shot^3^* mutant phenotype was significantly improved (Fig. 6G).

We then tested whether this increase in off-track MTs would correlate with more MT disorganisation. At 6 HIV, *shot^3^*mutant neurons displayed a 2.4-fold, and *Efa6^GX6[w-]^* mutant neurons a 1.55-fold increase in MDI (normalised to wild-type); this value was dramatically increased to 6.16 fold in *shot^3^ Efa6^GX6[w-]^* double mutant neurons (Fig. 8M). This strongly suggests that Efa6 and Shot do not act through the same mechanism, but perform complementary roles in regulating and maintaining axonal MTs and MT bundles. This conclusion was further confirmed by our finding that transfection of *Efa6-FL-GFP* into *shot^3^* mutant neurons could alleviate the MDI phenotype (Fig. 8N).

Finally, we assessed whether these complementary relationships between Shot and Efa6 are relevant *in vivo*. Since complete loss of Shot is an embryonically lethal condition, we first tested this in culture whether the lack of just one copy of *shot* has an enhancing effect on Efa6 deficiency. We found that MT disorganisation phenotypes of *Efa6-RNAi* (blue bar in Fig.8L) and of *shot^3/+^* heterozygous mutant neurons (orange bar) at 6 HIV were clearly enhanced when both genetic manipulations were combined (orange/blue bar). When testing the same genetic constellations in our optic lobe model, we found that the originally observed increase in MT disorganisation caused by cell-autonomous knock-down of *Efa6* (black arrows and blue bar in Fig.9B,E) was also further enhanced when the same experiment was carried out in a *shot^3/+^* heterozygous mutant background (black arrows and orange/blue bar in Fig.9C,E).

These findings support our conclusion that there is a correlation between off-track MTs and MT disorganisation. Furthermore, they are consistent with a scenario where both Shot and Efa6 regulate axonal MTs but through independent and complementary pathways: Efa6 inhibits MTs at the cortex (with peripheral MTs persisting for longer if Efa6 is absent), whereas Shot actively maintains MTs in bundles (with more MTs going off-track if Shot is absent) - and both these functions complement each other during MT bundle maintenance (see further details in the Discussion).

## Discussion

### Cortical collapse factors are important microtubule regulators relevant for axon morphology

Axons are the structures that wire our brain and body and are therefore fundamental to nervous system function. To understand how axons are formed during development, can be maintained in a plastic state thereafter, and why they deteriorate in pathological conditions, we need to improve our knowledge of axonal cell biology (Hahn et al., 2019). The MT bundles that form the core of axons are an essential aspect of this cell biology, and understanding how these bundles are regulated and contribute to axon morphogenesis will provide essential insights into axon development and maintenance (Voelzmann et al., 2016a). Here we have addressed fundamental contributions made by cortical collapse factors. We started from reports that two such factors from distinct protein families both negatively impact on axon growth in species as diverse as *C. elegans* (*Ce*Efa6; Chen et al., 2015; Chen et al., 2011) and mouse (Kif21A; van der Vaart et al., 2013).

We found that *Dm*Efa6 likewise acts as a negative regulator of axon growth. We demonstrate that fly Efa6 is a cortical collapse factor, inhibiting MTs primarily via the 18 aa long MTED. Since the MTED is the only shared motif with *Ce*Efa6 in an otherwise entirely divergent N-terminus (Fig. 3C), this clearly demonstrates that the MTED is functionally conserved between both species (Chen et al., 2015; Chen et al., 2011; O’Rourke et al., 2010).

Capitalising on *Drosophila* neurons as a conceptually well-established model (Prokop et al., 2013; Voelzmann et al., 2016a), we went on to demonstrate two novel roles for Efa6: as a negative regulator of axon branching and a quality control factor maintaining MT bundle organisation. To perform these functions, Efa6 does not affect the dynamics of MTs contained within the central axonal bundles, but it inhibits mainly those MTs that leave these bundles (Fig.10A). By inhibiting explorative MTs in GCs, it negatively impacts on a key event underlying axon growth (explained below; yellow arrows in Fig.10C). By inhibiting off-track MTs in the axon shaft, it tones down the machinery that seeds new interstitial branches (red arrow in Fig.10C), but also prevents these MTs from going astray and cause MT disorganisation (curled MTs in Fig.10C).

**Fig. 10.**
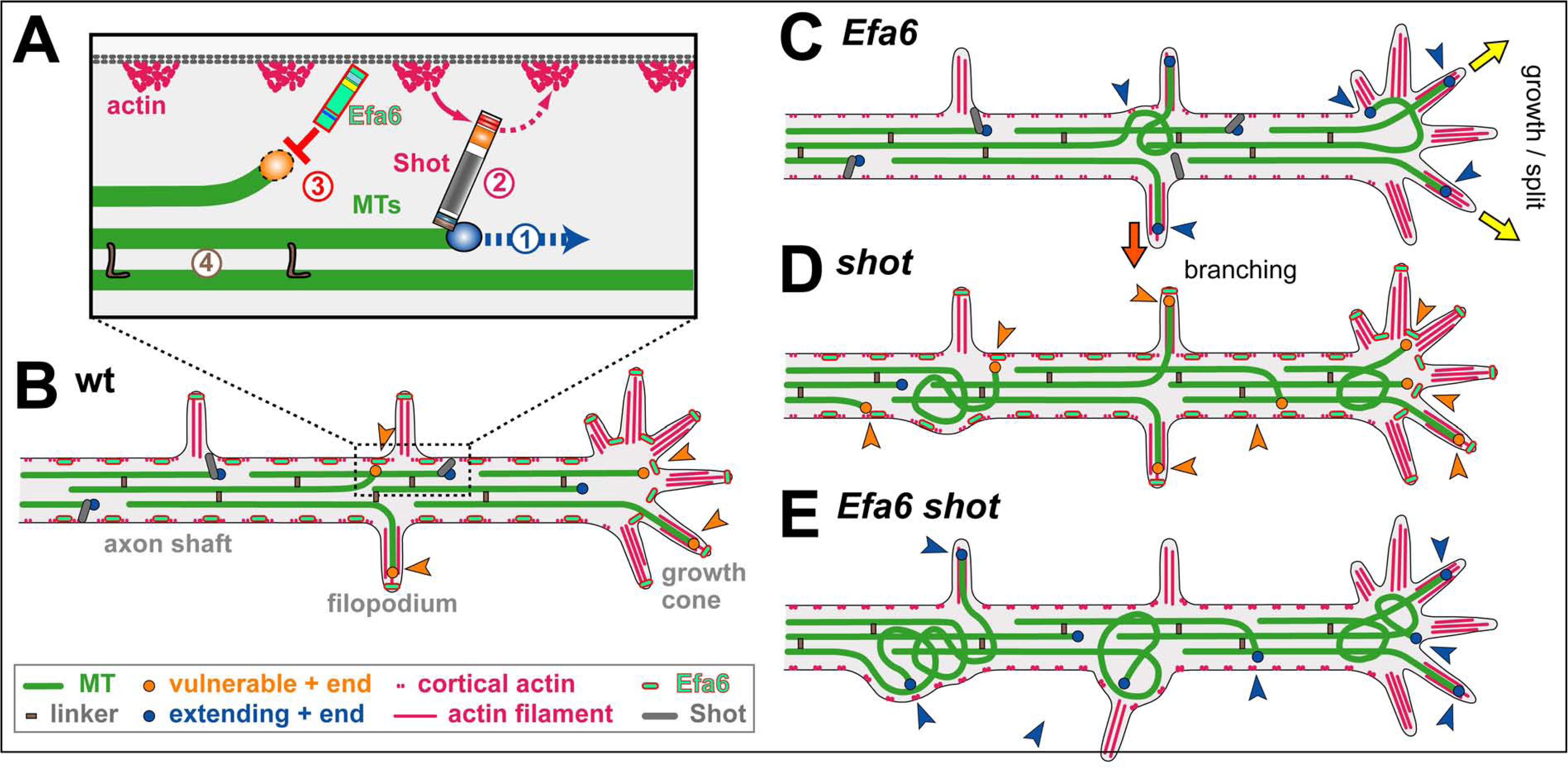
A model of the suggested roles of Efa6. **A**) The model of local axon homeostasis (Prokop, 2016; Voelzmann et al., 2016a) states that axonal MTs (green bars) are actively kept in bundles; for example, their continued polymerisation (1) mediated by plus end machinery (blue circle) is guided by spectraplakins (here Shot) along cortical actin into parallel bundles (2), or MTs are kept together through cross-linkage (brown “L”; 4) (Bettencourt da Cruz et al., 2005; Krieg et al., 2017); here we propose that MTs accidentally leaving the bundle become vulnerable (orange circles) through inhibition (red “T”) by cortically anchored Efa6. **B**) In normal neurons, polymerising MTs in bundles (dark blue circles) are protected by Shot (marine lines) and MT-MT cross-linkers (brown rectangles), whereas MTs approaches the membrane (orange arrow heads) either by splaying out in GCs or leaving the bundle in the shaft (orange arrow heads) become vulnerable (orange circles) to inhibition by Efa6 (light green/red dashes) both along the cortex and in filopodia **C**) Upon Efa6 deficiency, MTs leaving bundles or entering GCs are no longer subjected to Efa6-mediated cortical inhibition (blue arrow heads) and can cause gradual build-up of MT disorganisation; when entering shaft filopodia they can promote interstitial branch formation (red arrow), when entering GC filopodia they can promote axon growth or even branching through GC splitting (yellow arrows). **D**) Far more MTs leave the bundles in *shot* mutant neurons, but a good fraction of them is inhibited by Efa6 (increased number of orange arrow heads). **E**) In the absence of both Shot and Efa6, more MTs leave the bundles, but there is no compensating cortical inhibition (increased number of blue arrow heads), so that the MT disorganisation phenotype worsens.

Therefore, our work provides conceptual understanding of cortical collapse factors, which can explain how their molecular functions and subcellular roles in MT regulation link to their reported axonal growth phenotypes during development and regeneration (Chen et al., 2015; Chen et al., 2011; Heidary et al., 2008; van der Vaart et al., 2013), and to their additional functions in axon branching and maintenance reported here. Apart from existing links of cortical collapse factors to neurodevelopmental disorders (Heidary et al., 2008; Tiab et al., 2004; van der Vaart et al., 2013), we would therefore predict future links also to neurodegeneration.

### Roles of Efa6 during axonal growth

During axon growth, MTs constantly polymerise towards the periphery of GCs; the advance of many of these MTs is inhibited at the leading edge, and our work shows that cortical collapse factors are key mediators to this end. Only a fraction of MTs enters filopodia, potentially helped by active guidance mechanisms such as MT-actin cross-linkage (e.g. through spectraplakins, tau, drebrin-EB3; Alves-Silva et al., 2012; Biswas and Kalil, 2018; Geraldo et al., 2008). The widely accepted protrusion-engorgement-consolidation model of axon growth proposes that stabilised MTs in filopodia can seed axon elongation events (Aletta and Greene, 1988; Goldberg and Burmeister, 1986; Prokop et al., 2013). This model is consistent with our findings for Efa6. Thus loss of Efa6 can contribute to enhanced axon growth in two ways: firstly, through allowing more MTs to enter filopodia; secondly, by allowing them to dwell in filopodia for longer, thus enhancing the likelihood of their stabilisation (yellow arrows in Fig.10C). This scenario can explain why loss of *Efa6* in *C. elegans* improves axon re-growth after injury and growth overshoot during development (Chen et al., 2015; Chen et al., 2011), and why the upregulation of Kif21A levels in GCs causes stalled axon growth (van der Vaart et al., 2013).

### Roles of Efa6 during axonal branching

As explained in the introduction, axon branching can occur via GC split, in that diverging MTs get stabilised in parallel in the same GC (Acebes and Ferrus, 2000; yellow arrows in Fig.10C). Alternatively, it can occur through interstitial branching which involves the active generation (e.g. through MT severing) and then stabilisation of off-track MTs (Kalil and Dent, 2014; Lewis et al., 2013; Tymanskyj et al., 2017; Yu et al., 2008). Both models agree with our observations in Efa6-deficient/over-expressing neurons: we find greater/lower numbers of MTs in GC and shaft filopodia at 6 HIV, which then correlate with enhanced/reduced axonal branch numbers in mature neurons (red arrow in Fig.10C).

If interstitial branch formation is negatively regulated by *Efa6*, this poses the question as to whether Efa6 has to be actively down-regulated in healthy neurons for branching to occur; Efa6 could either be physically removed from future branch points (Chen et al., 2015) or its MT inhibition function could be switched off. We believe that no such regulation is required because Efa6 seems to be in a well-balanced equilibrium. Enough Efa6 appears to be present to inhibit occasional, likely accidental off-track MTs; this capacity is surpassed when the number of off-track MTs is actively increased, for example through MT severing proteins during axonal branch formation (Yu et al., 2008). Such a saturation model is supported by our experiments with *shot* (Fig.6F,G): filopodial MT numbers are elevated in *shot* mutant neurons, although Efa6 is present and functional (as demonstrated by the further increase in filopodial MT numbers in *shot Efa6* double-mutant neurons; Fig.10D,E). This strongly suggests that Efa6 function occurs at a level that is easily saturated when increasing the number of explorative MTs.

### Roles of Efa6 during axonal MT bundle maintenance

Axonal MT disorganisation in Efa6-deficient neurons occurs gradually and can even be induced by knock-down of Efa6 at mature stages (Fig.8K). Therefore, Efa6 appears to prevent MT disorganisation during axon development and maintenance, as is consistent with its continued expression in the nervous system (Fig.1). Such a continued role makes sense in a scenario where MT bundles remain highly dynamic throughout a neuron’s lifetime, constantly undergoing polymerisation to drive renewal processes that prevent senescence (Hahn et al., 2019; Voelzmann et al., 2016a).

Based on these findings, we propose Efa6 to act as a quality control or maintenance factor within our model of “local axon homeostasis” (Hahn et al., 2019; Voelzmann et al., 2016a). This model states that MTs in the force-enriched environment of axons have a tendency to go off-track and curl up (Pearce et al., 2018), thus potentially seeding MT disorganisation. Different classes of MT-binding regulators, amongst them spectraplakins, prevent this by actively promoting the bundled conformation (Voelzmann et al., 2017). We propose that cortical collapse factors act complementary in that they do not actively maintain MTs in bundles, but inhibit those MTs that have escaped the bundling mechanisms (Hahn et al., 2019).

In this scenario, MTs are protected from cortical collapse as long as they are actively maintained in axonal bundles; this can explain the long known conundrum of how axonal MTs extend hundreds of micrometres in relative proximity to the cell cortex in axons, whereas in non-neuronal cells cortical proximity of MTs tends to trigger either their inhibition or tethered stabilisation (Fukata et al., 2002; Kaverina et al., 1998).

### Evolutionary and mechanistic considerations of Efa6 function

We found that the MTED motif correlates well with MT inhibiting functions of Efa6 family members, whereas the rest of the N-terminus bears no obvious further reminiscence. Our experiments with N-terminal protein and synthetic MTED peptide, both reveal association with MTs/tubulin. The MTED strongly interferes with MT polymerisation. Future co-crystallisation experiments are required to reveal how the MTED works. Given its small size we hypothesise that it simply blocks assembly, rather than acting via more complex mechanisms such as active promotion of depolymerisation (e.g. kinesin-8 and -13, XMap215; Al-Bassam and Chang, 2011; Brouhard and Rice, 2014) or severing (e.g. spastin, katanin, fidgetin; McNally and Roll-Mecak, 2018; Sharp and Ross, 2012).

In any case, the small size of MTEDs might come in handy as experimental tools to inhibit MTs, potentially displaying complementary properties to existing genetic tools such as the kinesin-13 Kif2C (Moore et al., 2005; Schimizzi et al., 2010), stathmin (Marklund et al., 1996) or spastin (Eckert et al., 2012). Importantly, the experiments with the CAAX domain have shown that Efa6’s MT inhibiting function can be targeted to specific subcellular compartments clearing them of MTs, thus opening up a wide range of future applications.

Interestingly, the MT-inhibiting role of Efa6 seems not to be conserved in chordates (Fig.2A). However, roles of cortical collapse factors in neurons seem to have been taken over by other proteins such as the kinesin-4 family member Kif21A. The CFEOM1-linked Kif21A^R954W^ mutation causes the protein to relocate from the axon shaft to the growth cone of cultured hippocampal neurons (van der Vaart et al., 2013). In consequence, increased Kif21A levels in GCs cause reduced axon growth - and we observed the same with Efa6 over-expression (green bars in Fig.2D). The decreased levels of Kif21A in proximal axons correlate with a local increase in side branches - and we observed the same with Efa6 loss of function (blue bars in Fig.7E, L).

Finally, we found that the C-terminal domains of Efa6 might display some degree of functional conservation. So far, work on mammalian PSDs has revealed functions for C-terminal domains in regulating ARF6, ARF1 or ARL14 during actin cytoskeletal reorganisation and membrane ruffling, tumour formation and immune regulation (Derrien et al., 2002; Paul et al., 2011; Pils et al., 2005). Our finding that PSD1 and C-terminal Efa6 constructs cause similar membrane ruffling phenotypes in fibroblasts (Figs.S4 and S6), suggests that some conserved functions reside in this region and might further contribute, together with N-terminally mediated MT inhibition, to the neuronal or non-neuronal defects that cause semi-lethality displayed by *Efa6* mutant flies (data not shown).

### Conclusions and future perspectives

We propose that Efa6 acts as a cortical collapse factor which is important for the regulation of axonal MTs and relevant for axon growth, maintenance and branching. Although this function of Efa6 is evolutionarily not widely conserved, our findings provide a helpful paradigm for studies of other classes of cortical collapse factors also in mammalian neurons. Promising research avenues will be to refine our mechanistic understanding of how Efa6 blocks MT polymerisation, not only to better understand how it can be regulated in axons, but also to better exploit MTEDs as molecular tools in cell biological research.

## Acknowledgements

This work was made possible through support by the BBSRC to A.P (BB/I002448/1, BB/P020151/1, BB/L000717/1, BB/M007553/1) to N.S.S. (BB/M007456/1) and KD (BB/J005983/1), by parents as well as the Faculty of Life Sciences to Y.Q., by the Leverhulme Trust to I.H. (ECF-2017-247) and by the German Research Council (DFG) to A.V. (VO 2071/1-1). The Manchester Bioimaging Facility microscopes used in this study were purchased with grants from the BBSRC, The Wellcome Trust and The University of Manchester Strategic Fund. The Fly Facility has been supported by funds from The University of Manchester and the Wellcome Trust (087742/Z/08/Z). We thank Tom Millard and Marvin Bentley for very helpful comments on the manuscript, Simon Lowell for advice on the phylogenetic analyses, Hiro Ohkura for kindly providing DmEb1 antibody and Andrew Chisholm for the *C.elegans* Efa6 and human PSD constructs. Stocks obtained from the Bloomington *Drosophila* Stock Center (NIH P40OD018537) were used in this study.

## Supplementary materials

**Fig. S1.**
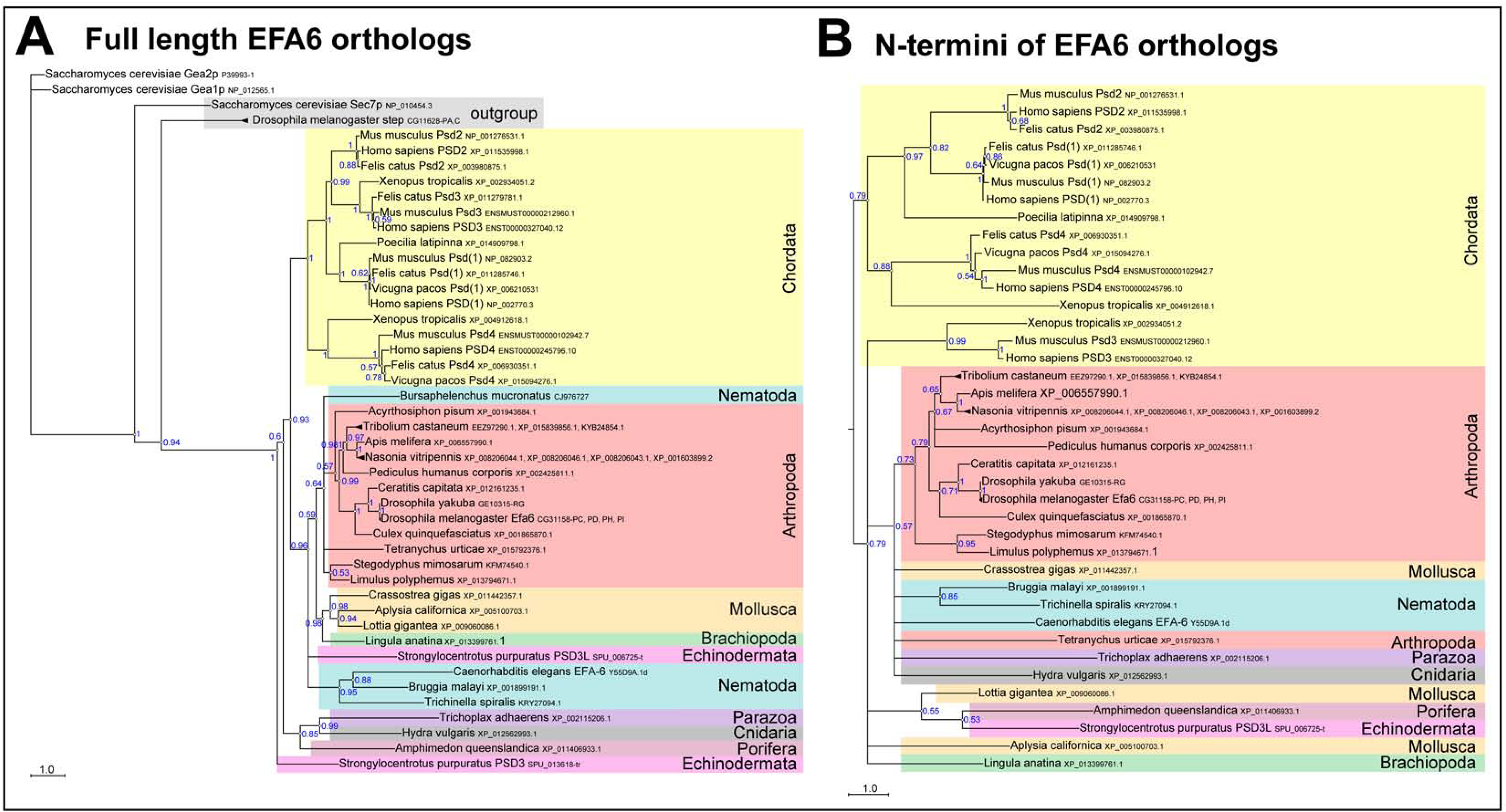
Phylogentic tree analysis of Efa6. Bayesian phylogenetic analysis of full length (A) or N-termini (B) of Efa6 orthologues. Sequences were aligned using Muscle or ClustalO and posterior probabilities and branch lengths calculated using MrBayes. Branch length scale is indicated; blue numbers show posterior probabilities of each branch split. For the full length tree, *Drosophila steppke* (*step*) was used as outgroup (Hahn et al., 2013). In both full length and N-terminus analyses, chordates (cream colour) split off very early from Efa6 versions of other species, in line with an early speciation event separating both groups before the vertebrate multiplication events took place. Phyla are highlighted in different colours, gene symbols and/or accession numbers are given after the species names.

**Fig. S2.**
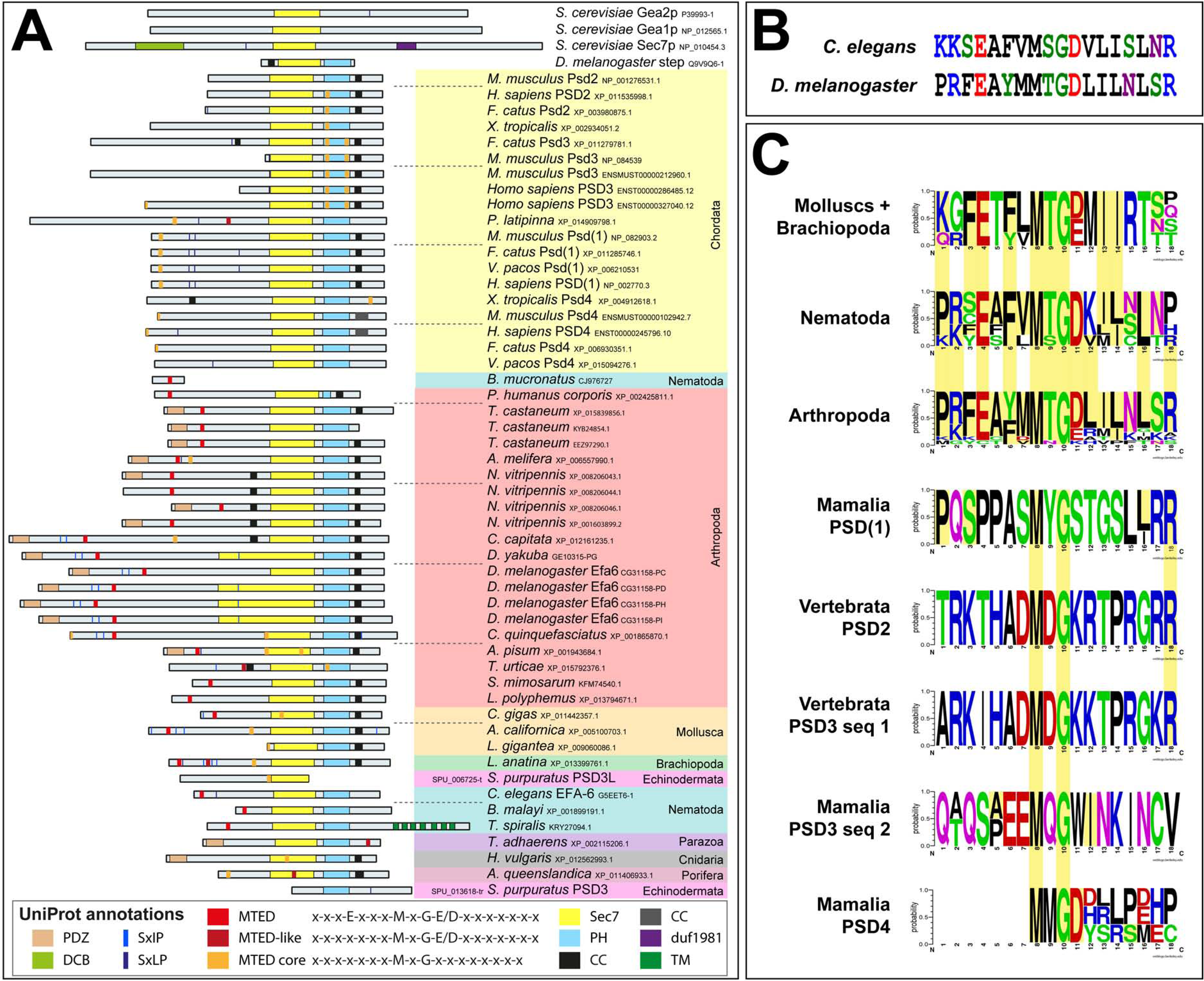
Efa6 N-terminal domains vary amongst different phyla. A) Domain annotations in 56 Efa6 orthologues via EMBL SMART and Uniprot. Phyla are colour-coded as in Fig.S1. Note that there is a strong variation of lengths and domain composition in particular of the N-terminus. The putative PDZ domain seems to be a feature mainly of insect versions of Efa6 and is absent from any analysed chordate orthologues; MTED and MTED-like sequences cannot be consistently identified in all Efa6 orthologues and are very divergent in chordate Efa6/PSD versions. SxIP/SxLP sites [flanked by positive charges as would be expected of functional motifs (Honnappa et al., 2009)] are found in the N-terminal half of only a subset of Efa6 versions in nematodes (e.g. *C. elegans*), insects (in particular flies, e.g. *D. melanogaster*) and molluscs, and even fewer in chordates; in mouse, alpaca and cat SxIP/SxLP sites are flanked by negative charges. B) To determine a potential MTED consensus sequence, 37 sequences of molluscs, nematodes, arthropods and putative MTED sequences of mammalian PSD1-4 were grouped according to phylum; consensus sequences were depicted using Berkley’s Weblogo online server (default colour scheme). Amino acid positions identical to *D. melanogaster* and *C. elegans* MTED are highlighted (faint yellow).

**Fig.S3.**
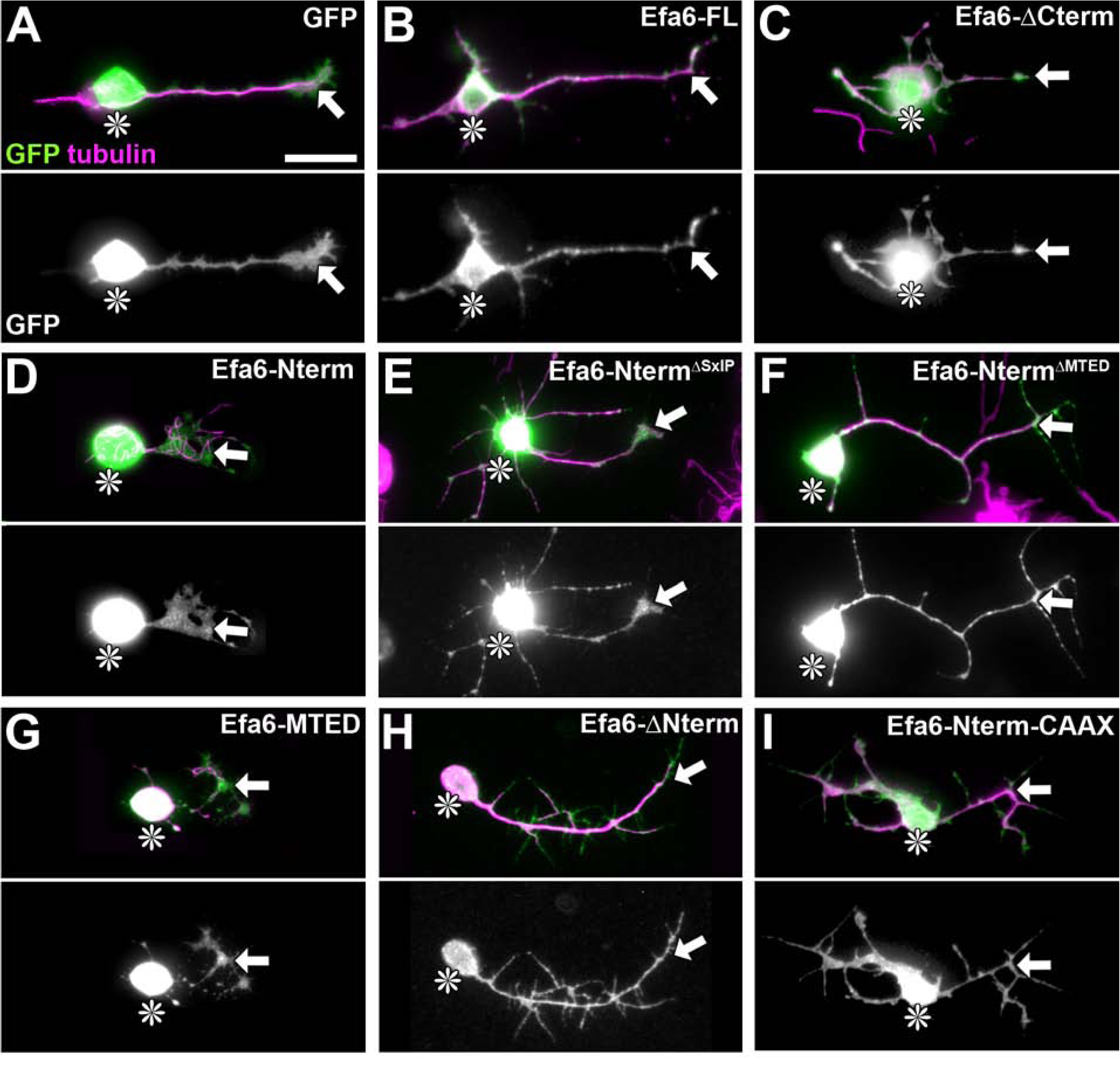
Localisation of Efa6 constructs in primary neurons. Images show transfected primary *Drosophila* neurons at 18HIV stained for tubulin (magenta) and GFP (green and in greyscale below the colour image). Cell bodies are indicated by asterisks, axon tips by arrows. The transfected constructs are indicated top right following the nomenclature explained in Fig.3B. Scale bar in A represents 10µm for all figures shown.

**Fig.S4.**
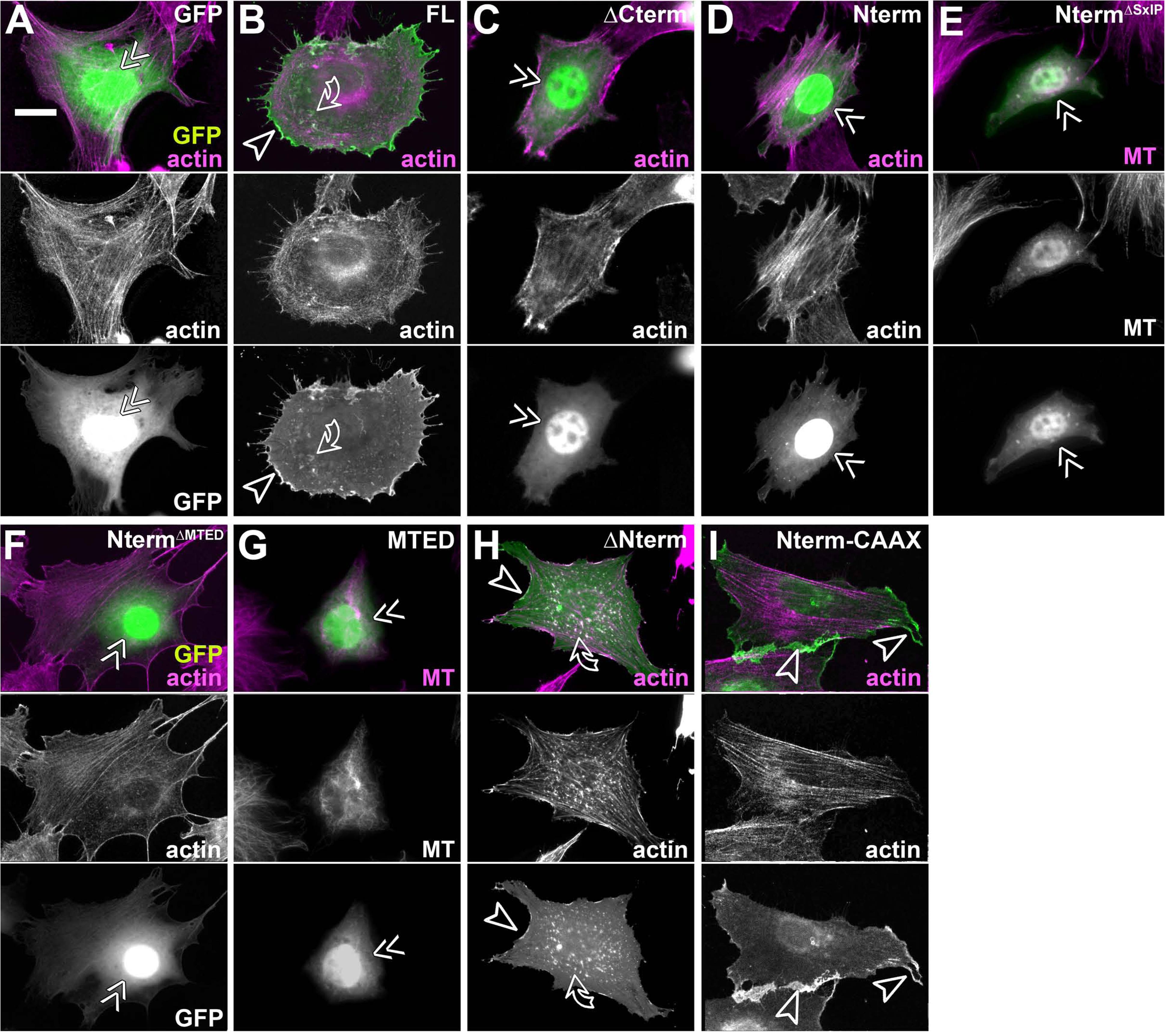
Efa6 constructs localisations in fibroblasts. Images show fibroblasts which are all stained for GFP (green) and either for actin or MTs (magenta); GFP and actin/MTs shown as single channels in greyscale below the colour image, 24hrs after transfection with control (GFP) or Efa6-derived constructs (nomenclature as explained in Fig.3B, but leaving out the “Efa6“-prefix and “::GFP”-postfix, as indicated top right. Double chevrons indicate nuclear localisation, arrow heads membrane localisation apparent at cell edges and curved arrows membrane ruffles. Scale bar in A represents 10µm in all images.

**Fig.S5.**
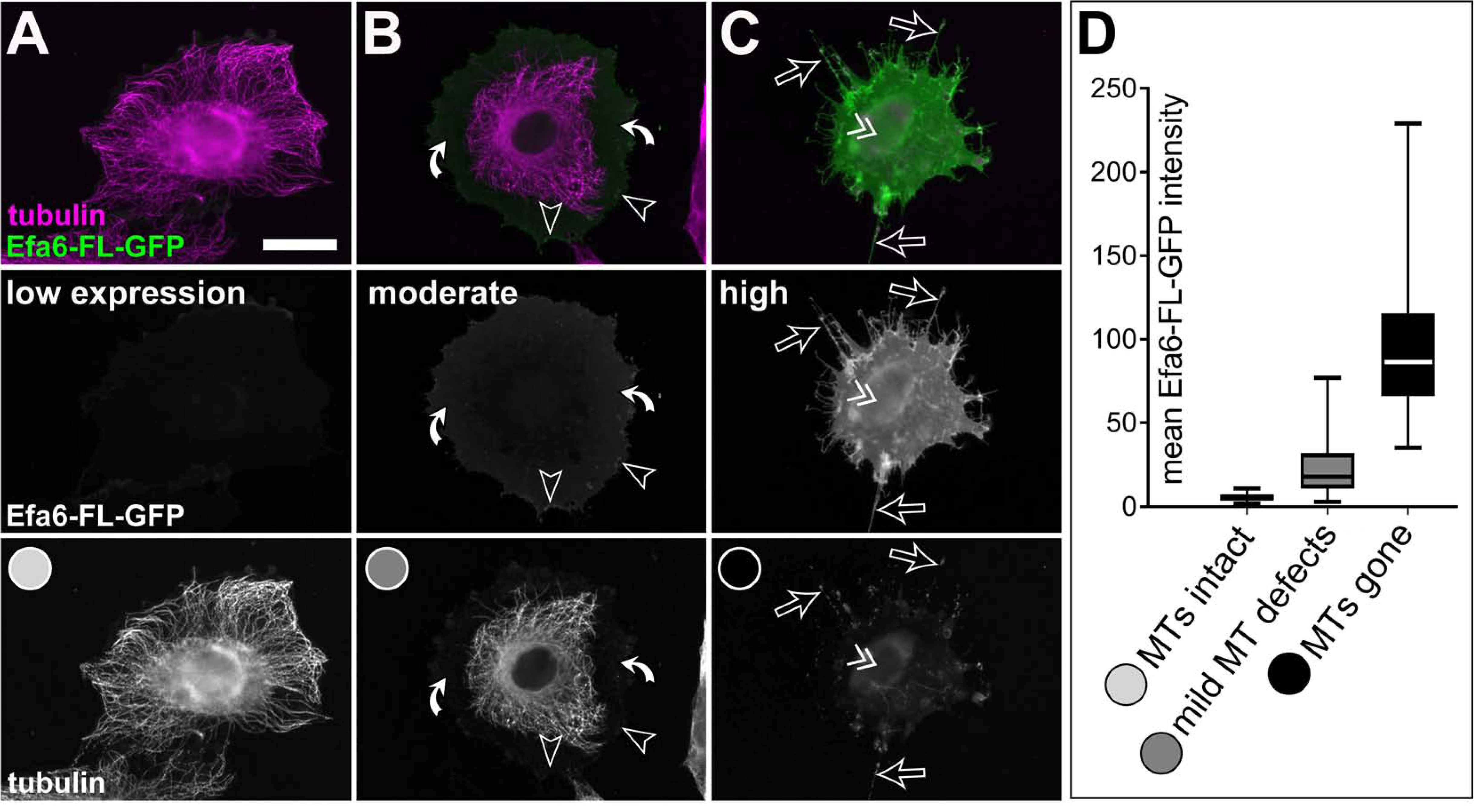
MT inhibition by Efa6-FL is concentration-dependent in fibroblasts. **A-C)** Representative images of fibroblasts stained for GFP and tubulin (green and magenta, respectively; both shown as single channels in greyscale below the colour image). Images were taken 24hrs after transfection with *Efa6-FL::GFP*, assessed for GFP intensity (plotted on the ordinate in **D**; examples for low, moderate and high expression are given in the left, middle and right columns, respectively) and then grouped with respect to their MT phenotypes into “MTs intact” (light grey), “mild MT defects” (medium grey) or “MTs gone” (black), as indicated by greyscale circles in the lowest row of A-C and the abscissa of D. Arrow heads point at GFP accumulation at membrane edges, white curved arrows indicate areas of the cell from where MTs have retracted, open arrows point at retraction fibres and the double chevron indicates the nucleus position and signs of nuclear GFP localisation. Scale bar in A represents 10µm in all images. For raw data see T9, which can be downloaded here: <u>w.prokop.co.uk/Qu+al/RawData.zip</u>.

**Fig. S6.**
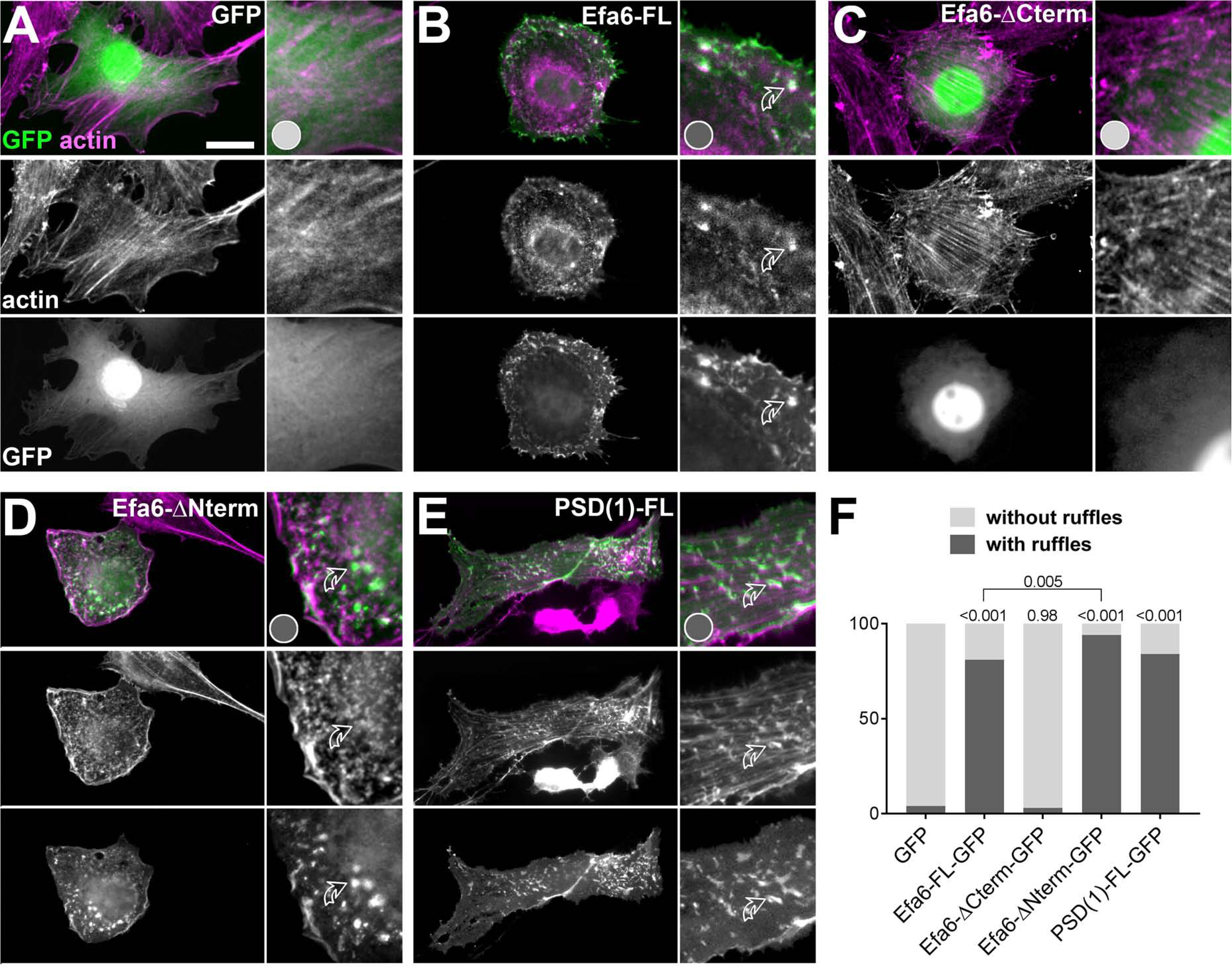
Conserved functions of the Efa6 C-terminus in membrane ruffle formation. **A-E)** Representative images of fibroblasts stained for GFP and actin (green and magenta, respectively; both shown as single channels in greyscale below the colour image). Images were taken 24hrs after transfection with different constructs (indicated top right): control vector (GFP; A), Efa6-derived constructs (B-D; nomenclature as explained in Fig.3B, but leaving out the “Efa6“-prefix and “::GFP”-postfix) or PSD1-FL::GFP (E). To the right of each image, a selected area is displayed with 2.5-fold magnification, showing dotted actin- and GFP-stained ruffles (curved open arrows) in B, D and E, but not A and C; ruffle formation has been quantified and is shown as a bar graph in **F**. All graph bars indicate percentages of fibroblasts with/without membrane ruffles (dark/light grey); P values on top of bars are from Chi^2^ tests relative to GFP controls. As shown, ruffle phenotype were never observed with any N-terminal Efa6 variants but are reproduced with the Efa6-ΔNterm::GFP variant (comprising the C-terminal Sec7, PH and CC domains; Fig. 3B). Scale bar in the top image of A represents 10 µM for all fibroblasts shown. For raw data see T10, which can be downloaded here: <u>w.prokop.co.uk/Qu+al/RawData.zip</u>.

**Fig. S7.**
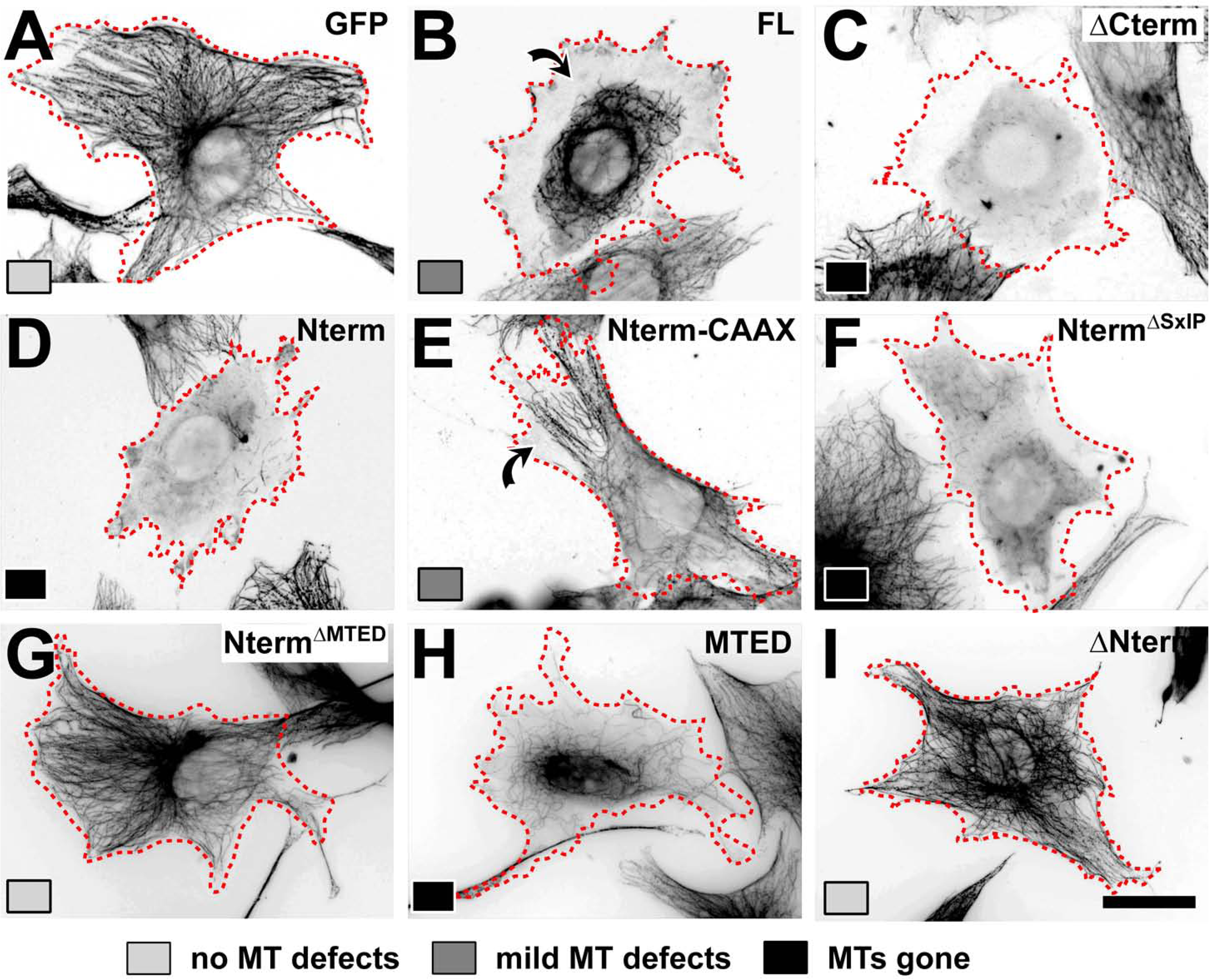
Representative MT phenotypes induced by the different constructs in transfected fibroblasts. Fibroblasts 24hrs after transfection with different control (GFP) or Efa6-derived constructs as indicated top right in each image (nomenclature as explained in Fig.3B, but leaving out the “Efa6“-prefix and “::GFP”-postfix). Cells were stained for tubulin (black; images shown as inverted greyscale) and classified as “no MT defects” (light grey), “mild MT defects” (medium grey) or “MTs gone” (black), as indicated by greyscale boxes bottom left of each image; curved arrows indicate peripheral MT depletion. Each image represents the most prominent phenotype for each respective construct. Scale bar at the bottom right in H represents 25µm in all images.

**Fig. S8.**
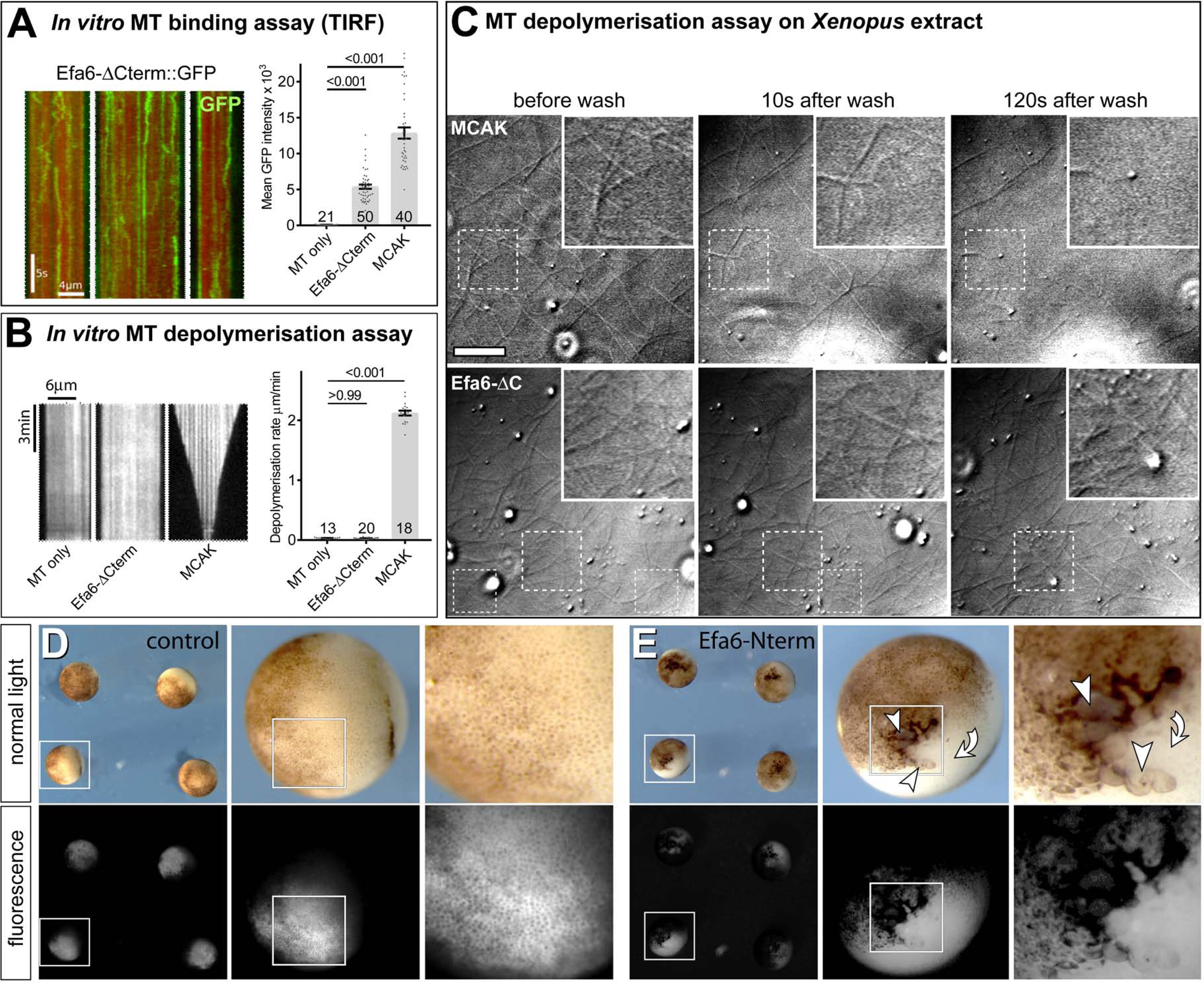
*In vitro* attempts to resolve the MT inhibition mechanism of Efa6. A) To determine whether Efa6 directly affects MT stability, we recombinantly expressed Efa6-ΔCterm::GFP in Sf9 cells, purified the protein and observed its interaction with MTs using total internal reflection fluorescence (TIRF) microscopy in a low ionic strength buffer (BRB20, 75mM KCl); images on the left show three examples of kymographs of MT lattices decorated with Efa6-ΔCterm::GFP, which displays a mixture of stationary molecules and diffusive interactions typical of non-translocating MT-associated proteins (Helenius et al., 2006; Hinrichs et al., 2012); bar charts (right) show quantification of the amount of interacting protein: background signal from MTs alone, Efa6-ΔCterm::GFP (20nM) and the non-translocating kinesin MCAK::GFP (20nM) as positive control; at the same protein concentration, over 2-fold more molecules of MKAC typically interact with MTs; all graphs in this figure show individual data points and bars representing mean ± SEM; numbers above bars show P values obtained from Mann–Whitney Rank Sum statistical analyses, numbers in bars the respective sample numbers. B) Kymographs (left) show individual fluorescently-labelled GMPCPP-stabilised MTs *in vitro* (Patel et al., 2016), either alone (MTs only; basal depolymerisation rate, n=7) or in the presence of 14 nM Efa6-ΔCterm::GFP (n=8) or 40nM MCAK::GFP (n=18); the bar chart (right) quantifies the induced depolymerisation rates; using two different purifications of Efa6-ΔCterm::GFP on three separate occasions, we saw no evidence of MT depolymerisation above the basal level of depolymerisation typically observed in these assays, whereas parallel control experiments with mitotic centromere-associated kinesin/MCAK showed MT depolymerisation rates typical of this kinesin. C) To assess whether MT destabilisation might require additional cytoplasmic factors, we used *Xenopus* oocyte extracts: phase contrast images show MTs in *Xenopus* oocyte extracts (after they had been allowed to polymerise for 20 min) and then showing stills from before, 10s after and 120s after washing in 20nM MCAK::GFP (as positive control) or 20nM Efa6-ΔCterm::GFP; squares outlined by dashed white lines are shown magnified in insets revealing MTs; MTs clearly vanish upon treatment with MCAK, but counts of MTs did not reveal any obvious effects on MTs with Efa6-ΔCterm::GFP. D,E) *RFP* controls and *Efa6-Nterm::GFP* expression constructs were injected into *Xenopus* embryos at the 4-cell stage and analysed 24 hrs later; only the *Efa6* construct caused a strong suppression of cell division, as indicated by the presence of very large cells (arrows) and pigmentation defects (curved arrows) at the site of injection, suggesting that Efa6-Nterm::GFP is functional when expressed in the *Xenopus* context. Scale bar (in top left image) represents 3 µm in C (2.5 fold magnified insets), and 1400 µm / 350 µm / 140 µm in left / middle / right images of D and E, respectively. For raw data see T11, which can be downloaded here: <u>w.prokop.co.uk/Qu+al/RawData.zip</u>.

**Fig. S9.**
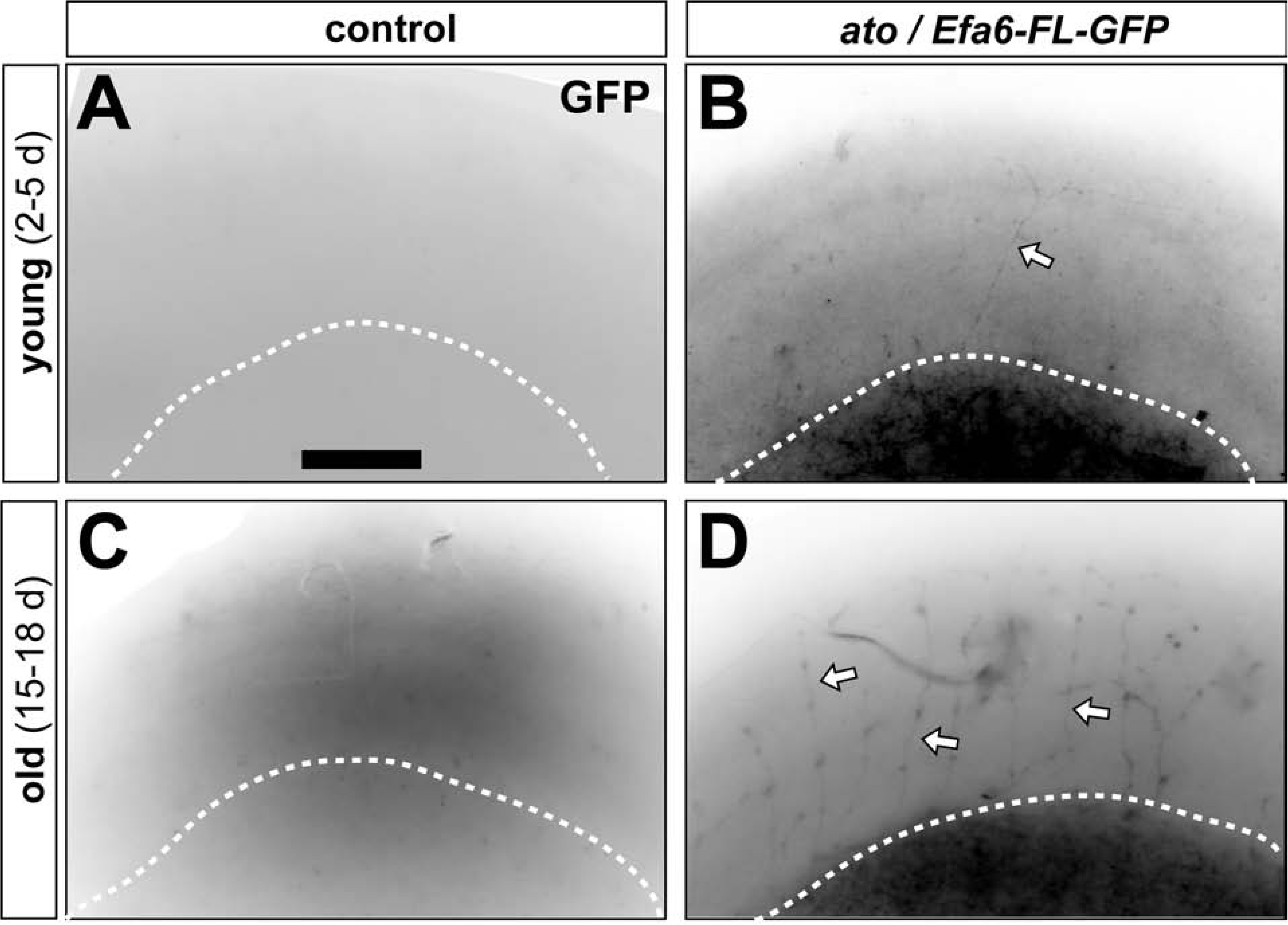
*ato-Gal4*-driven Efa-FL::GFP expression in adult brains. Brains (region of the optic lobe in oblique view; white dashed line indicating the lower edge of the medulla) of young (2-5 d after eclosure; top) and old flies (15-18 d; bottom) which are either from wild-type controls or from flies driving *UAS-myr-tdTomato* and *UAS-Efa6-FL-GFP* via the *ato-Gal4* driver in dorsal cluster neurons; specimens are stained for GFP and arrows point at stained; for myr-dtTomato staining of brains from these experiments see Fig.7F-K. Scale bar in D represents 60µm in all images.

**Fig. S10.**
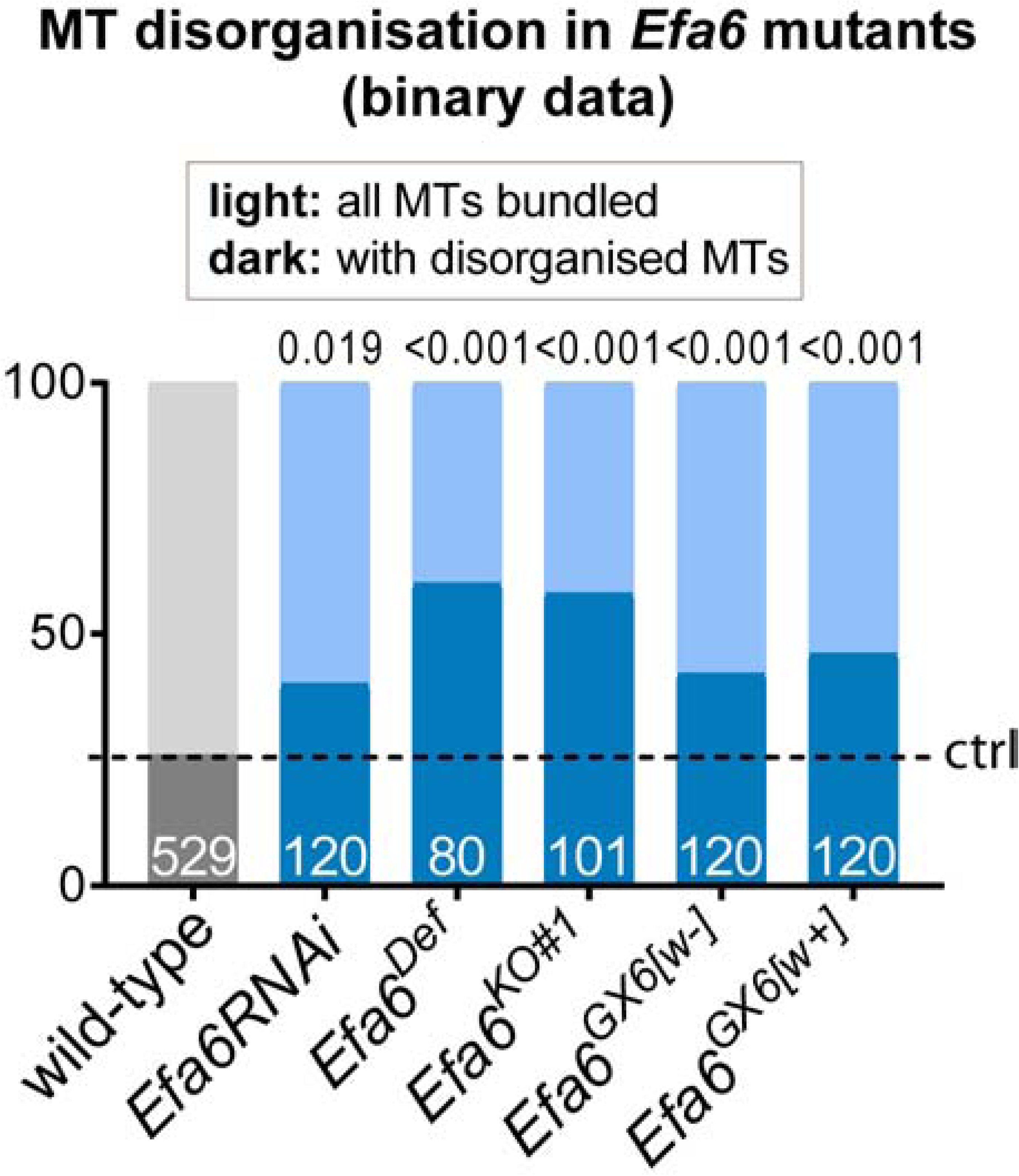
Efa6 mutant primary neurons show MT disorganisation. Quantitative analyses of MT disorganisation measured as binary score: percentages of neurons with bundled MTs throughout (light colour) or showing obvious MT disorganisation in their axons (dark colour) in different *Efa6* mutant alleles (control value indicated as horizontal dashed “ctrl” line); P values from Chi^2^ tests are given above the bars. For raw data see T12, which can be downloaded here: <u>w.prokop.co.uk/Qu+al/RawData.zip</u>.

**Suppl. Mov. M1**. MT behaviours in control fibroblasts transfected with GFP. Eb3 (green) was used to visualise MT polymerisation, GFP (magenta) localises throughout the cell; MTs clearly polymerise to the very fringe of the transfected cell. Movies were taken at one frame per second; the scale bar corresponds to 25μm, close ups are twofold magnified. Movie available at <u>w.prokop.co.uk/Qu+al/SupplMov.html</u>.

**Suppl. Mov. M2**. MT behaviours in fibroblasts transfected with Efa6-FL::GFP. Eb3 (green) was used to visualise MT polymerisation, which is prevented from entering areas of high Efa6-FL::GFP (magenta), with Efa6 often accumulating where MTs enter the area. Movies were taken at one frame per second; the scale bar corresponds to 25μm, close ups are twofold magnified. Movie available at <u>w.prokop.co.uk/Qu+al/SupplMov.html</u>.

**Suppl. Mov. M3**. Eb1::GFP in mouse NHTH3 fibroblast about 6 hours after transfection. Pictures for the movie were taken at 2 s intervals. Scale bar indicates 10 µm. Movie available at <u>w.prokop.co.uk/Qu+al/SupplMov.html</u>.

**Suppl. Mov. M4**. MT behaviours in fibroblasts transfected with Efa6-Nterm::GFP::CAAX. Eb3 (green) was used to visualise MT polymerisation, which is prevented from entering areas of high Efa6-FL::GFP (magenta), with Efa6 often accumulating where MTs enter the area. Movies were taken at one frame per second; the scale bar corresponds to 25μm, close ups are twofold magnified. Movie available at <u>w.prokop.co.uk/Qu+al/SupplMov.html</u>.

**Suppl. Mov. M5**. Eb1::GFP in a growth cone of a wild-type *Drosophila* primary neuron at 6 HIV. Arrows indicate positions where individual Eb1::GFP comets terminate. Pictures of the movie were taken at 2 s intervals. Scale bar indicates 10 µm. Movie available at <u>w.prokop.co.uk/Qu+al/SupplMov.html</u>.

**Suppl. Mov. M6**. Eb1::GFP in a growth cone of a wild-type *Drosophila* primary neuron at 6 HIV. Arrows indicate positions where individual Eb1::GFP comets terminate. Pictures of the movie were taken at 2 s intervals. Scale bar indicates 10 µm. Movie available at <u>w.prokop.co.uk/Qu+al/SupplMov.html</u>.

**Suppl. Mov. M7.** Eb1::GFP in a growth cone of an *Efa6 ^GX6[w-]^* mutant *Drosophila* primary neuron at 6 HIV. The arrow heads follow individual Eb1::GFP comets illustrating either their trajectories adjacent to the membrane or prolonged dwell time at filopodial tips. Pictures for the movie were taken at 2 s intervals. Scale bar indicates 10 µm. Movie available at <u>w.prokop.co.uk/Qu+al/SupplMov.html</u>.

**Suppl. Mov. M8.** Eb1::GFP in a growth cone of an *Efa6 ^GX6[w-]^* mutant *Drosophila* primary neuron at 6 HIV. The arrow heads follow individual Eb1::GFP comets illustrating either their trajectories adjacent to the membrane or prolonged dwell time at filopodial tips. Pictures for the movie were taken at 2 s intervals. Scale bar indicates 10 µm. Movie available at <u>w.prokop.co.uk/Qu+al/SupplMov.html</u>.

## Notes

#### Summary of Updates

This version contains important new experiments: (1) New in vitro data show that the MTED binds tubulin and blocks its polymerisation in the absence of other proteins; (2) live imaging and transfected fibroblasts shows that areas enriched with Efa6-FL or Efa6-Nterm::CAAX prevent MTs from elongation; (3) transfection of shot mutant neurons with Efa6-FL rescues their MT disorganisation phenotypes.

http://www.prokop.co.uk/Qu+al/RawData.zip

http://www.prokop.co.uk/Qu+al/SupplMov.html

